# Photocaged chloroquine derivatives for the light-dependent inhibition of autophagy in cancer stem cells

**DOI:** 10.1101/2025.11.24.690081

**Authors:** Sofía Alonso-Manresa, Carme Serra, Lourdes Muñoz, Marina Bataller, Yoelsis Garcia-Mayea, Matilde Esther Lleonart Pajarin, Belen Garcia Prats, Sandra Mancilla Zamora, Zamira Vanessa Diaz Riascos, Amadeu Llebaria, Laia Josa-Culleré

**Author notes:** Corresponding authors: Amadeu Llebaria, and Laia Josa-Culleré,.

## Abstract

Chloroquine (CQ) and hydroxychloroquine (HCQ) inhibit autophagy and have shown promise as adjuvant anticancer agents, particularly for targeting therapy-resistant cancer stem cells (CSCs). However, their clinical utility is limited by systemic toxicity and poor tumour selectivity. Here we report the design, synthesis, and photochemical evaluation of [7-(diethylamino)coumarin-4-yl]methyl (DEACM)-caged CQ and HCQ derivatives as visible-light-activated autophagy inhibitors. Selective caging of the aliphatic amine fully suppressed biological activity in the dark and enabled rapid, efficient release of the parent drugs upon illumination. The lead compound **1C** displayed robust light-dependent cytotoxicity across multiple cancer cell lines and, upon photoactivation, recapitulated CQ’s effects on LC3-II accumulation. In CSC-enriched tumourspheres, illumination of **1C** completely abolished spheroid formation, demonstrating precise spatiotemporal control of stemness suppression. Importantly, *ex vivo* and *in vivo* studies confirmed that visible light penetrates tumour tissue sufficiently to activate **1C** and release CQ within the tumour. These findings establish the first proof of concept for light-controlled autophagy inhibition and provide a blueprint for spatiotemporally confined anticancer therapies based on photopharmacological modulation of CSCs.

## Introduction

Chloroquine (CQ) and its derivative hydroxychloroquine (HCQ), initially developed and widely used as antimalarials,^1^ were later repurposed and approved for the treatment of autoimmune disorders such as rheumatoid arthritis and systemic lupus erythematosus.^2^ They have also been explored as antivirals, including for HIV-1, SARS, and MERS.^3–5^ More recently, numerous *in vitro* and *in vivo* studies have investigated the potential of CQ and HCQ for oncology, either as single agents or in combination with cytotoxic regiments.^6–8^ Several clinical trials are currently ongoing (https://clinicaltrials.gov/), mainly focused on evaluating their potential as adjuvant therapy combined with chemotherapeutic drugs and radiotherapy.^9–11^

As examples, HCQ is being evaluated in combination with the CD4/6 inhibitor abemaciclib to target residual disease in breast cancer^12^ and with Akt inhibitor MK2206 for advanced solid tumours.^13^

Compelling evidence indicates that CQ and HCQ can target cancer stem cells (CSCs) – a minor, highly tumorigenic subpopulation that drives therapeutic resistance and relapse.^14,15^ CSCs exhibit stem-like properties (self-renewal, differentiation), enhanced drug resistance, and long quiescent phases that render them less susceptible to antiproliferative agents.^16,17^ Once the bulk cancer cells are eliminated through chemotherapy, CSCs can differentiate into bulk cancer cells and regenerate the tumour. CSC enrichment has been documented in residual disease and metastases across cancer types,^18^ including head and neck squamous cell carcinoma (HNSCC)^19^ and triple-negative breast cancer (TNBC).^20^ Therefore, developing effective therapies that specifically target treatment-resistant CSCs may prevent local recurrences and distant metastases, improving overall survival and patient outcomes.^21^

Several *in vitro* and *in vivo* studies support the ability of CQ/HCQ to target CSCs, alone or in combination.^22,23^ As examples, CQ sensitises liver CSCs to the tumour microenvironment,^24^ targets CSCs in TNBC,^25,26^ potentiates temozolomide cytotoxicity in glioma cells,^27^ and, in combination with the tyrosine kinase inhibitor afatinib and with cisplatin^14^ eradicates CSCs of HNSCC.^28^ Clinical trials are also evaluating the ability of HCQ to overcome resistance in combination with chemotherapy,^29,30^ including taxanes,^31^ vorinostat,^32^ temozolomide,^33^ or bortezomib,^34^ and radiation therapy.^35^

While the precise anticancer mechanisms of CQ/HCQ remain incompletely defined, inhibition of autophagy is widely recognised as a principal mode of action. CQ accumulates preferentially in lysosomes, elevating intralysosomal pH, size, and permeability owing to its diprotic weak-base character (pKa of 8.1 and 10.2).^36^ Unprotonated CQ diffuses freely across membranes, but once inside acidic organelles such as the lysosome, it becomes protonated and trapped,^37^ leading to lysosomal alkalinisation and dysfunction that compromise proteolysis, chemotaxis, phagocytosis, and antigen presentation.^38^ In turn, lysosomal disfunction sensitises cancer cells towards cell death. Raising lysosomal pH also inhibits the fusion between autophagosomes and lysosomes, thereby impairing lysosomal protein degradation and autophagy.^39^

In addition, the basic centres of CQ/HCQ underlie other effects relevant to cancer therapy,^9^ including cytosolic acidification that perturbs aerobic metabolism and helps overcome chemoresistance,^40,41^ and Golgi de-acidification that alters glycosylation – another cancer hallmark. ^42,43^

Even though CQ and hydroxychloroquine HCQ show promise as adjunct therapies in oncology, and specifically in targeting the resistant CSC population, their clinical use in prolonged treatment regimens – such as those required for cancer – is limited by toxicity. Clinical studies have reported adverse effects including retinal toxicity, neuromyopathy, and lysosomal storage disorders that can progress to cardiomyopathy.^44–47^ HCQ is generally better tolerated than CQ and can be dosed higher in humans – daily doses of CQ are safe up to 500 mg, while HCQ is dosed up to 1200 mg/day.^48^ For this reason, the majority of clinical trials employ HCQ for combination therapy.^49^ Nevertheless, high-dose HCQ can still cause adverse events (fatigue, anorexia, gastrointestinal effects),^33^ and in some examples, these doses produced only modest autophagy inhibition *in vivo*,^50^ and dose-limiting toxicity can preclude escalation to pharmacodynamically effective exposures.^35^

In such cases where toxicity limits the use of drugs, photopharmacology emerges as a promising solution.^51,52^ It is based on the use of light to control the biological effect of drugs in a precise place and time. In oncology, it could allow us to restrict their effect to the tumour area only, avoiding undesired effects in other tissues. Rendering freely diffusible drugs responsive to light relies on two possible designs, giving photoswitches and photocages. Photoswitches contain a photoresponsive functional group, such as an azobenzene^53^ or a hemithioindigo,^54^ that isomerise under irradiation of a particular wavelength and intensity. The differences in polarity and geometry of the two isomers can result in a change in target affinity and efficacy, providing a reversible control of biological activity. The design is often based on modifying the structure of known drugs with the photoswitching moiety. Instead, for photocages the original drug is modified at a position that is essential for its biological activity with a photocleavable moiety, such as an *o*-nitrobenzyl or a coumarin.^55^

Numerous CQ/HCQ analogues have been described, which enhance lysosomal accumulation and cytotoxicity through structural tuning. As representative examples, the 4-alkyl chain has been substituted with a cymantrene group^56^ and longer aliphatic chains,^57^ the primary alcohol of HCQ has been used to introduce larger substituents and conjugates,^58^ and the 7-Cl has also been modified.^57^ Yet, to our knowledge, photopharmacology has not been applied to spatiotemporally control the effect of autophagy inhibitors. Given that the therapeutic use of CQ/HCQ is limited by a lack of tumour selectivity, we hypothesised that introducing a light-responsive moiety could enhance its use. Here we describe the design, synthesis, and photochemical characterisation of coumarin-caged CQ and HCQ derivatives and evaluate their light-dependent activity in breast and HNSCC models, including CSC-enriched tumourspheres.

## Results

### Design and synthesis of photocaged (H)CQ derivatives

Converting a known drug onto a light-responsive molecule often relies on introducing a photocaging moiety onto a functional group that is essential for its activity. Given the role of the basic amine(s) on the mechanism of CQ, we reasoned that protecting its basic centre(s) with a photocage could abolish its biological activity, which would be recovered upon uncaging under irradiation. We prepared derivatives of both CQ and HCQ to compare photochemical behaviour and biological responses, noting HCQ’s improved tolerability yet reports of greater CQ efficacy in some contexts.^48^ While we did not expect significant differences in their photochemical properties, we were interested in comparing their biological properties. Initial attempts with *o*-nitrobenzyl as the photocaging moiety led to low release yields and rates (data not shown); hence we present herein our results with [7-(diethylamino)coumarin-4-yl]methyl (DEACM) as the photocaging moiety.

Direct reaction of commercial chloroquine diphosphate with Br-DEACM resulted in low conversion to the protected product (**Scheme 1**). Instead, upon prior neutralisation with NaOH,^59^ CQ readily reacted with Br-DEACM to give a mixture of three major protected derivatives. CQ reacted at the tertiary aliphatic amine to give quaternary salt **1C**, at the 4-aniline to give tertiary amine **2C**, and at both sites to give the doubly protected product **3C**. These were separated by reverse-phase column chromatography and isolated as their formate salts. Regiochemical assignments for **1C** vs **2C** derived from Heteronuclear Multiple Bond Correlation (HMBC) experiments, which showed correlations between the methylene bridge of coumarin and the quinoline skeleton for **2C**, while for **1C** the correlations of the benzylic position of coumarin were with the methylene groups adjacent to the aliphatic quaternary amine (**Figure S1**).

**Scheme 1.**
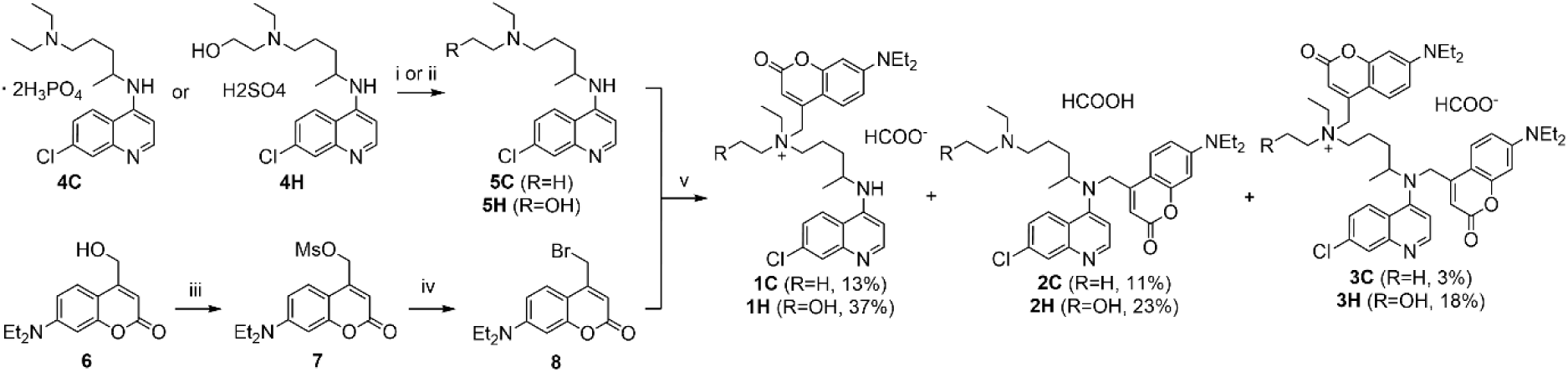
Synthesis of caged derivatives. i. for **5C**: NaOH, water, rt, 30 min, quant.; ii. for **5H**: NH_4_OH, CHCl_3_, rt, 1 h, quant.; iii. MsCl, Et_3_N, DCM, rt, 2 h, quant.; iv. LiBr, THF, rt, 18 h, quant.; v. MeCN, 60 °C, 18 h.

A similar strategy led to the three photocaged derivatives of HCQ **1H**,**2H**,**3H**.

### Photochemical characterisation of photocaged derivatives

With the six photocages in hand, we proceeded to evaluate their ability to release CQ/HCQ under irradiation (**Figure 1**). Under our standard HPLC methods, CQ appeared as a broad band, likely due to its basic nature. To enable its quantification, a method for the simultaneous HPLC-MS/MS analysis of CQ, HCQ, and their cage derivatives was developed, evaluating different sample solvents, stationary and mobile phases. Optimal chromatographic performance was achieved using a C18 stationary phase, 50 mM ammonium formate at pH3 as the mobile phase, and samples were prepared in a water-acetonitrile mixture containing at least 90% water. The use of the buffered mobile phase and using predominantly water to dissolve samples improved peak symmetry and sharpness by stabilising the ionisation state of CQ and minimising secondary interactions with the stationary phase.

**Figure 1.**
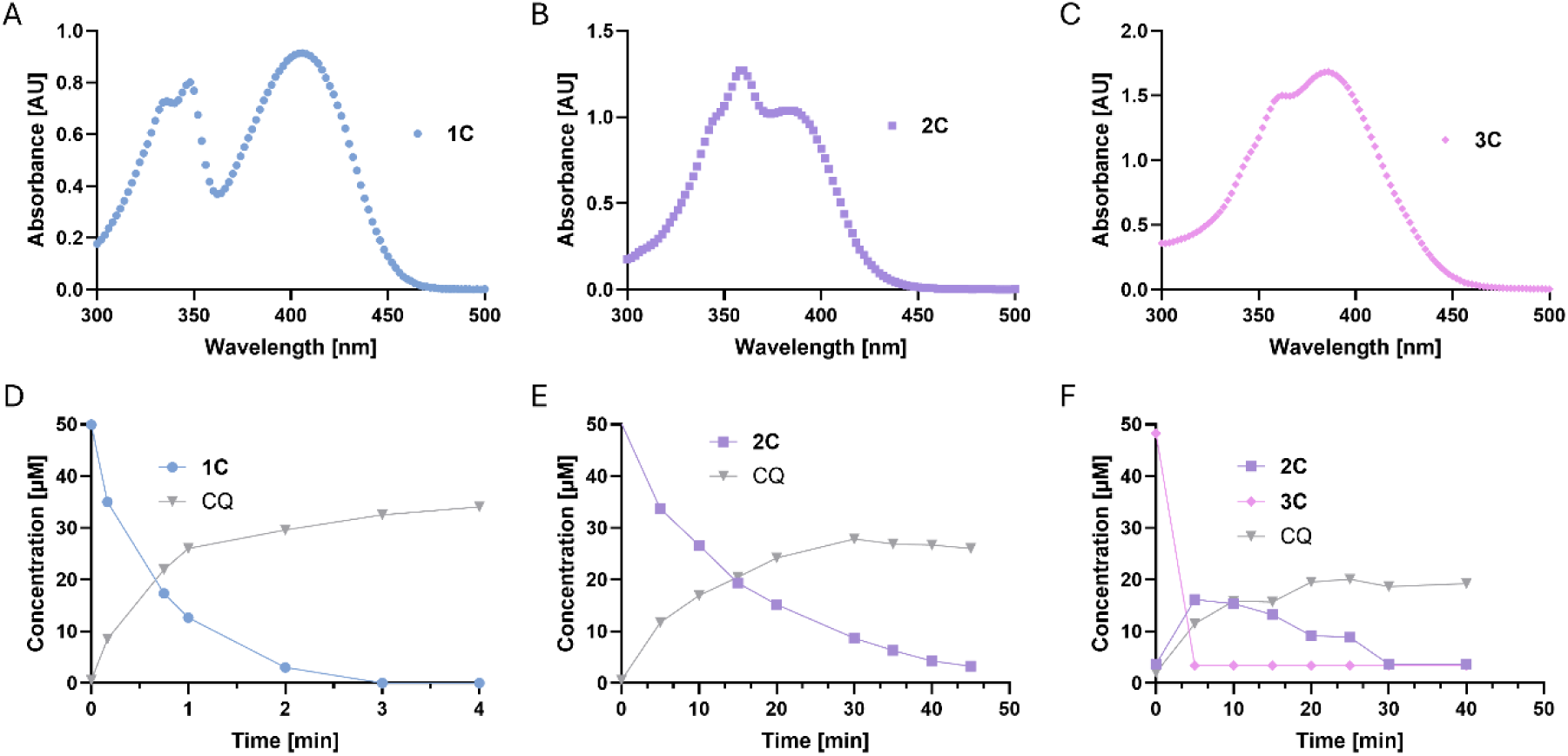
**Photochemical characterisation of compounds 1C-3C**. (A-C) Absorbance spectra of compounds **1C**-**3C**, at 100 µM in PBS. (D-F) HPLC quantification of photocage and CQ during illumination at 405 (**2C** and **3C**, 19 mW/cm^2^) or 420 nm (**1C**, 13 mW/cm^2^) for the indicated times, initial solution at 50 µM in PBS.

Firstly, we confirmed that the six compounds were stable in the dark under the conditions of the biological studies. After 3 days in complete cell culture medium at 37 °C, there were no significant changes in the concentration of the photocages, confirming that the active (H)CQ will not be released under lack of illumination (**Figure S2**).

Upon illumination, the coumarin at the aliphatic amine cleaved faster than at the aniline: **1C** released CQ with t₉₀ ≈ 3 min, whereas **2C** required 20 min under matched conditions (**Scheme 1**). The yield was also higher for **1C**. Consistently, double protected analogue **3C** released one coumarin rapidly to give **2C**, which then uncaged more slowly, at an overall lower yield of 40%. The isolated CQ recovery for **1C** (68%) aligns with reported yields for coumarin PPGs.^60,61^ HCQ derivatives behaved similarly: **1H** released HCQ with t₉₀ ≈ 2 min and a maximal concentration of 28 µM (57% yield) (**Table 1**, **Figure S3**).

**Table 1.** Photophysical and photochemical properties for the uncaging of **1C-3C** and **1H-3H**.

| Compound | $\lambda_{\text{max}}$ [nm]<sup>a</sup> | $\epsilon_{405}$ [M<sup>-1</sup>cm<sup>-1</sup>] | $\epsilon_{420}$ [M<sup>-1</sup>cm<sup>-1</sup>] | $t_{90}$ [min]<sup>b</sup> | Yield<sup>c</sup> |
| --- | --- | --- | --- | --- | --- |
| 1C | 406 | 9100 | 7900 | 3 | 68% |
| 2C | 360, 384 | 6600 | 2300 | 21 | 56% |
| 3C | 386 | 12900 | 7400 | 20 | 40% |
| 1H | 404 | 6700 | 5600 | 2 | 57% |
| 2H | 358, 384 | 5900 | 1800 | 35 | 51% |
| 3H | 388 | 17000 | 10200 | 10 | 24% |

<sup>a</sup>Wavelength at which absorption is maximum; <sup>b</sup>time required to reach 90% of the final concentration of (H)CQ; <sup>c</sup>chemical yield of released (H)CQ measured by HPLC, after illumination at 405 nm (2C and 3C, 19 mW/cm<sup>2</sup>) or 420 nm (1C, 13 mW/cm<sup>2</sup>). In all cases, 100 μM in DMSO.

We also quantified the release of coumarin alcohol **6** during the photolysis of **1H** and found that **6** only reached a concentration of 5 µM that remained stable after 2 min (**Figure S3D**), suggesting the formation of other photolytic products of the coumarin skeleton that might degrade, as we did not detect other discernible peaks in the HPLC spectra (**Figure 2C**).

**Figure 2.**
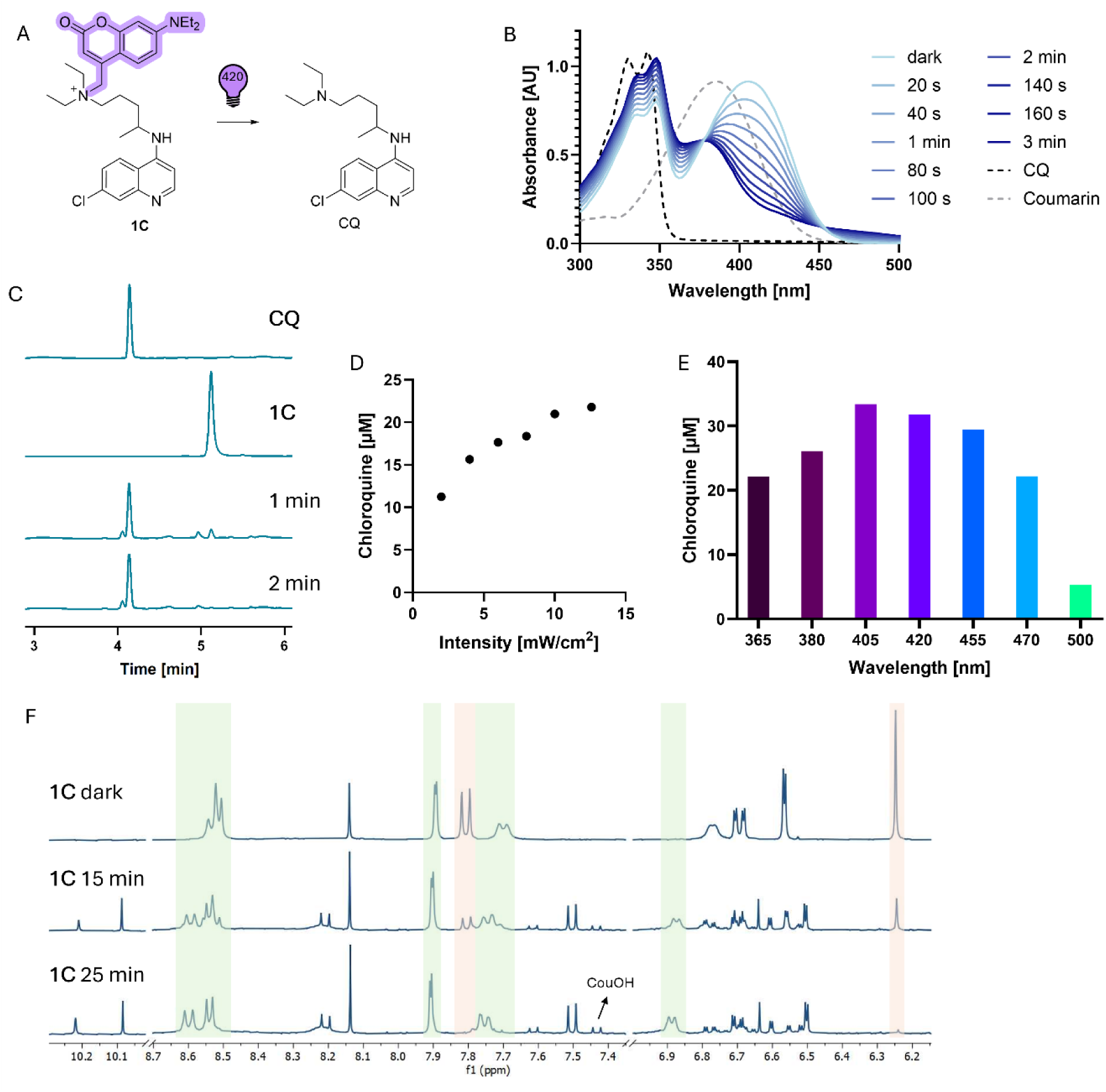
**Uncaging profile of 1C**. A) Uncaging reaction. B) Changes of the UV-Vis absorption of **1C** recorded in 20-second intervals during 420 nm (13 mW/cm^2^) irradiation at 100 µM in PBS, dotted lines show the UV-Vis absorption of CQ and coumarin **6** at 100 µM in PBS. C) Representative HPLC traces of CQ, **1C**, and **1C** after 420 nm (13 mW/cm^2^) illumination (1 and 2 min) at 100 µM in PBS. D) Amount of CQ formed after illumination of **1C** (50 µM in PBS) for 2 minutes at 420 nm of different intensities, quantified by HPLC. E) Amount of CQ formed after illumination of **1C** (50 µM in PBS) for 2 minutes at 365, 380, 405, 420, 455, 470, and 500 nm (11-13 mW/cm^2^ except for 365 at 7 mW/cm^2^), quantified by HPLC. F) ^1^H NMR of **1C** under dark and illumination (420 nm, 1 and 2 min), 2 mg/mL in DMSO-*d*_6_, green regions indicate peaks from the CQ quinoline scaffold and orange from the coumarin scaffold.

We selected the fastest photocage **1C** (**Figure 2A**) for a deeper photochemical analysis. Following the photolysis process by UV-Vis spectroscopy confirmed that the uncaging reaction is completed after 3 minutes at 420 nm with loss of the coumarin band and retention of characteristic CQ band (**Figure 2B**); a similar profile was observed for **1H** (**Figure S4A**). This is consistent with the photolysis studies by HPLC (**Figure 2C**), which had shown CQ as the major released product. We also followed the uncaging process by ^1^H NMR (**Figure 2F**), which also showed disappearance of the coumarin peaks with minimal changes on CQ quinoline signals. In this experiment, the photolysis required a longer time to complete as the amount and concentration of photocage was larger (100 µM for HPLC studies, 4 mM for NMR studies).

Illumination of **1C** for 2 minutes at 420 nm of different intensities (2–13 mW/cm²) gave an intensity-dependent release of CQ (**Figure 2D**), consistent with a photon flux-dependent reaction rate.^62^ Testing the photolysis at different wavelengths ranging from UV to green light (**Figure 2E**) indicated that, while 405 and 420 nm led to the fastest release consistent with its maximum absorbance at 406 nm (**Table 1**), similar conversions are obtained with up to 470 nm. Although at 500 nm the photolysis reaction was significantly slower, the release was still considerable with only 2 minutes of illumination giving a 10% release.

### Biological evaluation *in vitro*

Having confirmed that the prepared photocages are able to release (H)CQ under illumination, we tested their ability to induce a light-dependent decrease in viability of cancer cell lines. We first verified dark inactivity.

As breast cancer is one of the malignancies where autophagy inhibition by (H)CQ has been validated, we started our studies with the common breast cancer cell line MCF7. Cages bearing coumarin at the aromatic amine (mono- (**2C**, **2H**) or di- (**3C**, **3H**) substituted) retained appreciable activity at the highest concentration, approaching the IC_50_ of free (H)CQ (**Figure 3A, Figure S5**), and were therefore unsuitable as silent prodrugs. In contrast, the quaternised aliphatic-amine cages were minimally active in the dark: **1C** was inactive up to 200 µM and **1H** showed only weak effects at the top dose.

**Figure 3.**
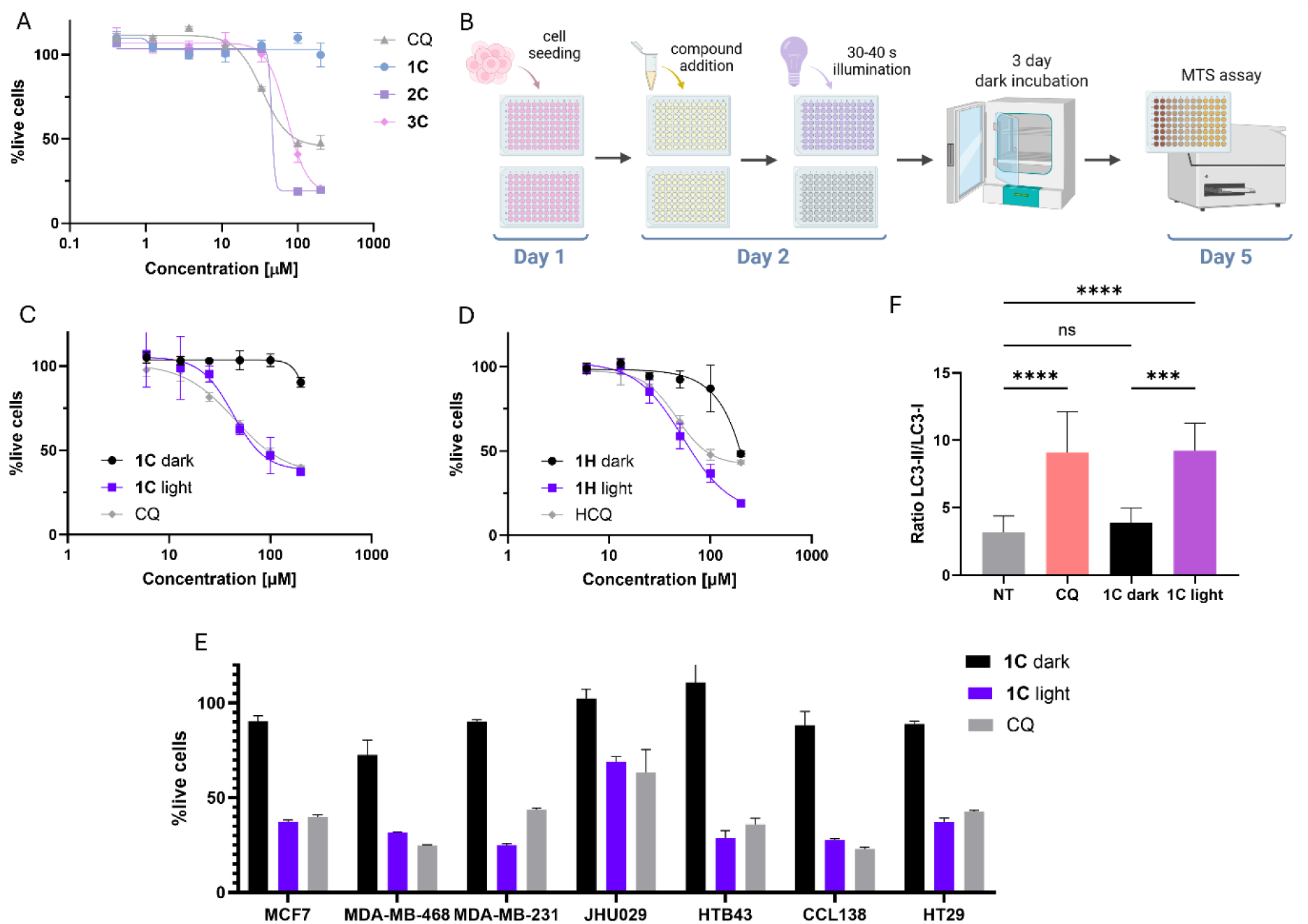
**Effects of photocages in cancer cell lines on adherent culture**. A) Dose-response curves of CQ and of the CQ photocages **1C-3C** on the viability of MCF7 cells (MTS assay) under dark conditions for 72 h (IC_50_ (CQ) = 34.1 µM; IC_50_ (**2C**) = 45.6 µM; IC_50_ (**3C**) = 72.6 µM). B) Workflow for viability studies with adherent cells under dark vs illumination conditions. C) Dose-response curves on the viability of MCF7 cells (MTS assay) of **1C** under dark vs illumination (420 nm, 8 mW/cm^2^, 40 s) conditions and of CQ for 72 h. D) Dose-response curves on the viability of MCF7 cells (MTS assay) of **1H** under dark vs illumination (420 nm, 8 mW/cm^2^, 30 s) conditions and of HCQ for 72 h. E) Viability of a range of cell lines upon treatment with **1C** under dark vs illumination (420 nm, 8 mW/cm^2^, 40 s) conditions and of CQ at 200 µM. F) Immunoblotting analysis on the effect of **1C** under dark vs illumination (420 nm, 8 mW/cm^2^, 1 min) conditions and of CQ on the relative expression levels of LC3-II and LC3-I on MCF7 cells; cells treated for 2 h. Values in A), C), D), E), and F) are represented as mean ± SD.

Other important controls were confirming that the cells are not affected by illumination at 420 nm for 40 s to 2 min (**Figure S6**) and that the activity of CQ is not affected by illumination (**Figure S7**).

Short illumination immediately after dosing (**Figure 3B**) restored activity: **1C** and **1H** displayed dose-dependent cytotoxicity matching their parent drugs over 72 h (**Figure 3C,D**). This light-dependent effect was reproduced across a range of cell lines covering breast cancer (MCF7, MDA-MB-468, MDA-MB-231), HNSCC (JHU029, HTB43, CCL138), and colon cancer (HT29) (**Figure 3E**, **Table 2**). In general, we observed a larger window between dark and light conditions for **1C** than **1H**, as **1H** showed some activity at the highest concentration(s).

**Table 2.** Effect of selected compounds on the viability of a range of cell lines under dark or illuminated conditions, measured by an MTS assay.

| Cell line | CQ<sup>a</sup> | HCQ<sup>a</sup> | **1C** |  | **1H** |  |
| --- | --- | --- | --- | --- | --- | --- |
|  |  |  | dark | light<sup>b</sup> | dark | light<sup>b</sup> |
| MCF7 | 44.6 | 46.5 | >200 | 42.6 | ≈200<sup>c</sup> | 53.5 |
| MDA-MB-231 | 35.2 | 39.7 | >200 | 51.5 | ≈200 | 41.5 |
| MDA-MB-468 | 57.4 | 55.4 | na<sup>d</sup> | 69.9 | ≈200 | 46.6 |
| JHU029 | ≈200 | 91.2 | na | ≈200 | ≈200 | 85.8 |
| HTB43 | 102.1 | 98.2 | na | 107.4 | na | 97.7 |

<sup>a</sup>Cells kept in the dark only.
<sup>b</sup>Cells illuminated at 420 nm (8 mW/cm<sup>2</sup>) for 30-40 s before the 72-hour incubation period.
<sup>c</sup>Estimation, full dose-response not obtained.
<sup>d</sup>na = not active, indicating no decrease in viability observed up to 200 μM.

To gain further evidence on the effect of **1C** being triggered through the release of CQ, we assessed by Western blotting for LC3-II protein, a common marker of autophagy.^63^ We observed that **1C** under dark conditions had similar levels of LC3-II as the non-treated cells, while **1C** under illumination triggered an increase on LC3-II accumulation similar to CQ (**Figure 3F**, **Figure S8**).

Coumarin alcohol **6**, which we had determined to be the major by-product of the uncaging reaction and was stable to the illumination conditions employed (**Figure S9**), showed no activity by itself in cellular viability (**Figure S10**). Nonetheless, we did observe that at longer illumination times and higher intensities, **6** showed some activity by itself and the photocages gave an increase in potency, suggesting that under these conditions the photocage could be acting through an alternative mechanism alongside autophagy inhibition.

One of the promises of autophagy inhibitors such as CQ is their effect on CSCs, hence, we also assessed the light-dependent effect of photocage **1C** on tumourspheres. Sphere cultivation is widely used to enrich CSCs from bulk cancer cells and is widely accepted as a functional assay of self-renewal property of CSCs.^64–66^ It is based on plating a single cell suspension at a proper cell density on ultralow attachment surface with the serum-free culture medium in supplementation with several defined growth factors. We grew MCF7, JHU029, and HTB43 cells under non-adherent conditions to form 3D spheres and confirmed these to be enriched in stemness factors (**Figure 4A,B**).^14^

**Figure 4.**
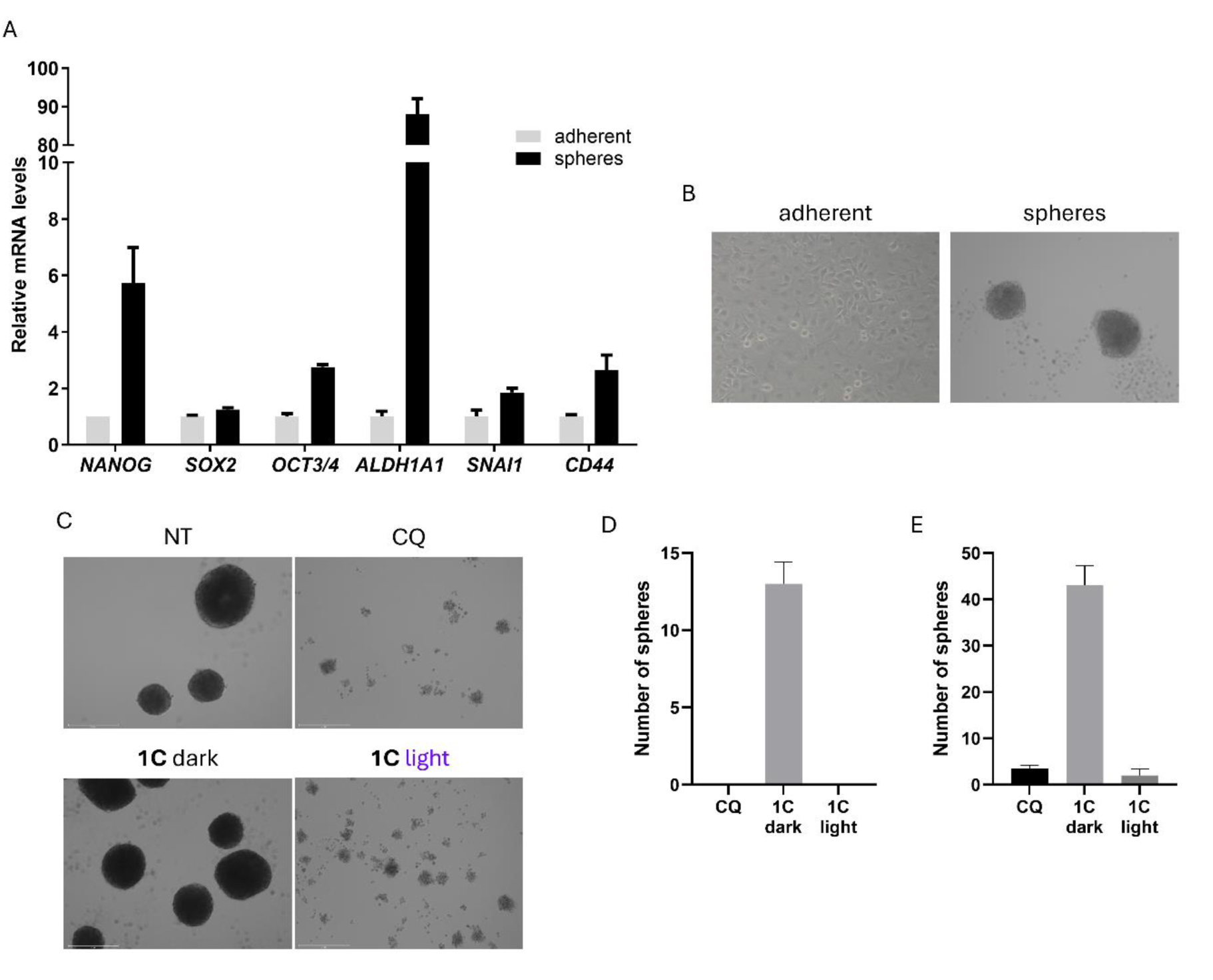
**Effect of 1C on CSC spheres**. A) mRNA levels of stemness genes in JHU029 cells grown in either adherent or sphere (first generation) culture, determined by PCR. B) Brightfield images of JHU029 cells grown in either adherent or sphere (first generation) culture. C) Brightfield images of HTB43 cells grown in sphere culture, non-treated or treated with **1C** at 100 µM under dark vs illumination (420 nm, 8 mW/cm^2^, 40 s) conditions and of with CQ at 100 µM for 10 days. Average number of spheres counted on individual wells of 96-well plates for JHU029 (D) and HTB43 (E) cells grown in sphere culture, treated with **1C** at 100 µM under dark vs illumination (420 nm, 8 mW/cm^2^, 40 s) conditions and of CQ for 10 days. Values in A), D), and E) are represented as mean ± SD.

To evaluate compound effects on CSCs, cells were treated immediately after seeding, and both sphere formation and viability were assessed after 7–10 days, when untreated spheres reached an average diameter of ≥250 µm for HTB43 and ≥300 µm for JHU029. Spheres were markedly more sensitive to CQ than adherent cultures (e.g., JHU029: IC₅₀ adherent ≈ 200 µM vs spheres ≈ 12 µM), consistent with autophagy’s role in CSC survival. As we did not observe major changes on the viability of spheres of first or third generation treated with HCQ (**Figure S11**), we continued the subsequent studies with first generation spheres.

Cells treated with **1C** and kept in the dark successfully formed spheres. In contrast, a brief 40-second illumination prior to the 10-day spheroid culture completely abolished their ability to form spheres in both JHU029 and HTB43 cells (**Figure 4C-E**, **Figure S12**). Illuminated **1C** exhibited lower IC₅₀ values compared to those obtained in viability assays in adherent cells, further supporting its impact on autophagy and CSCs (**Table 3**).

**Table 3.**
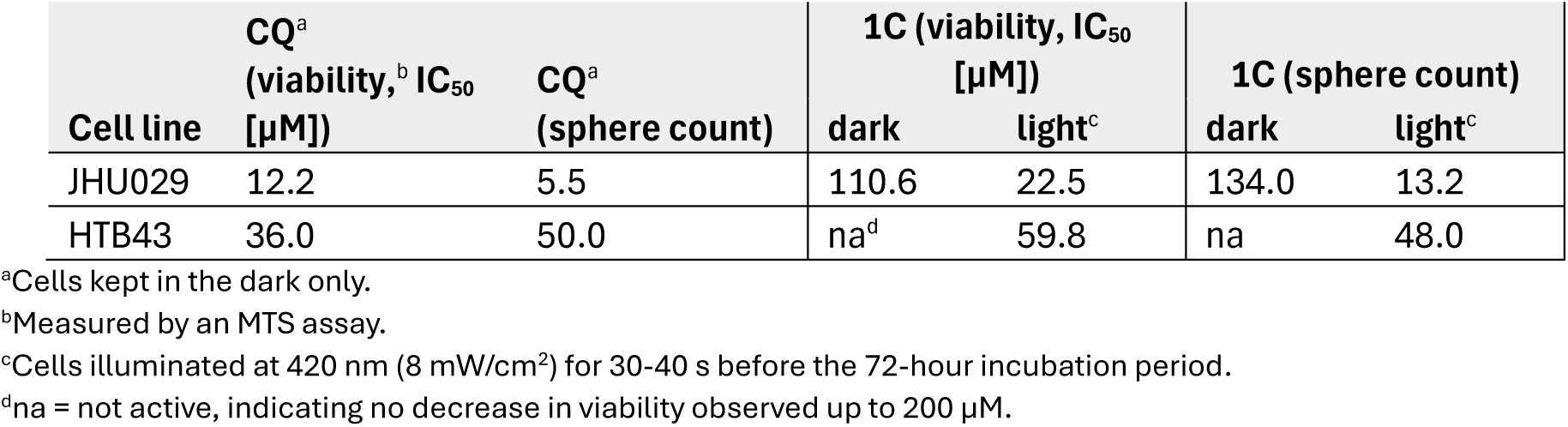
Effect of CQ and **1C** on CSC spheres of JHU029 and HTB43 cells under dark or illuminated conditions.

| Cell line | CQ<sup>a</sup><br>(viability, <sup>b</sup> IC<sub>50</sub><br>[μM]) | CQ<sup>a</sup><br>(sphere count) | 1C (viability, IC<sub>50</sub><br>[μM]) |  | 1C (sphere count) |  |
| --- | --- | --- | --- | --- | --- | --- |
|  |  |  | dark | light<sup>c</sup> | dark | light<sup>c</sup> |
| JHU029 | 12.2 | 5.5 | 110.6 | 22.5 | 134.0 | 13.2 |
| HTB43 | 36.0 | 50.0 | na<sup>d</sup> | 59.8 | na | 48.0 |

<sup>a</sup>Cells kept in the dark only.
<sup>b</sup>Measured by an MTS assay.
<sup>c</sup>Cells illuminated at 420 nm (8 mW/cm<sup>2</sup>) for 30-40 s before the 72-hour incubation period.
<sup>d</sup>na = not active, indicating no decrease in viability observed up to 200 μM.

### Photorelease *in vivo*

Having confirmed the fast release of photocage **1C** under illumination and its light-dependent effect in cellular studies *in vitro*, we assessed its ability to release CQ *in vivo*; we chose MDA-MB-231 tumours as this cell line gave a big window between dark and illuminated conditions.

We first quantified the amount of light transmitted through *ex vivo* mouse breast tumour samples of varying widths. Three wavelengths were selected – 405, 430, and 470 nm (**Figure S13**) – corresponding to the maximum uncaging efficiency of **1C** (**Figure 2E**). Illumination was performed using mounted LEDs from Thorlabs equipped with collimators to minimise scattering, and transmitted irradiance was measured with a photodiode power sensor (**Figure S14**). As expected, thicker tumours generally allowed less light to pass through (**Figure S15A,B**). However, this relationship was more heterogeneous compared to more homogeneous tissues such as chicken breast (**Figure S15C,D**), where attenuation correlated perfectly (r=0.976) with thickness following a logarithmic relationship. Across all samples, transmitted irradiance increased with wavelength: 470 nm > 430 nm > 405 nm (**Figure 5A**), consistent with reduced absorption at longer wavelengths. At full LED power (130 mW/cm² at 470 nm), up to 20 mW/cm² crossed the tumours – approximately 30% of the incident light. These results support that light can reach these tumours at a sufficient photon quantity to activate photocage **1C**, which we proceeded to confirm in a real animal setting.

**Figure 5.**
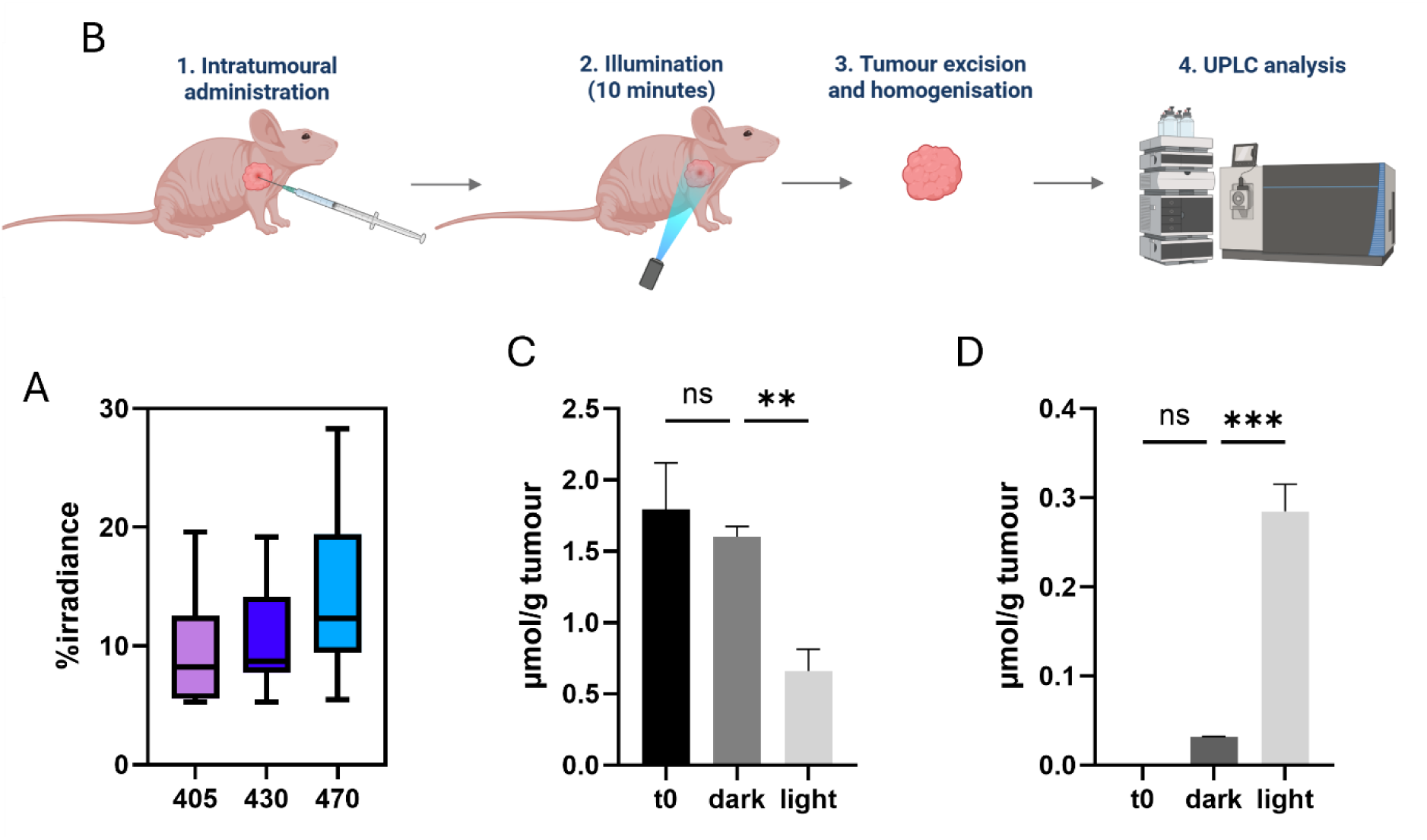
**Photolysis studies *ex vivo* and *in vivo.*** (A) Percentage irradiance detected after illumination of tumour samples of MDA-MB-231 cells *ex vivo* at the indicated wavelengths. (B) Workflow followed for the *in vivo* study, aimed at quantifying the amount of CQ released in an orthotopic breast cancer model with MDA-MB-231 cells. (C) Quantification of **1C** detected by HPLC before and after illumination of the tumour at 470 nm for 10 min, using the setup shown in (B). (D) Quantification of CQ detected by HPLC before and after illumination of the tumour at 470 nm for 10 min, using the setup shown in (B). Values in A), C), and D) are represented as mean ± SD.

As 470 nm resulted in the largest intratumour irradiance amongst the three tested wavelengths, and we confirmed that this light can be used to induce the same light-dependent effect of **1C** (**Figure S16**) as the previously used 420 nm, we performed the *in vivo* study with this wavelength (**Figure 5B**). We chose an orthotopic breast cancer model of MDA-MB-231 cells; cells were inoculated at the intramammary fat pad, and the study was initiated when tumours reached 80-120 mm^3^. We injected **1C** intratumorally at 10 mg/kg, and after illumination for 10 minutes, tumours were excised, homogenised, and analysed by HPLC. We found that the 10-minute illumination period was sufficient to consume 50% of the photocage (**Figure 5C**), with a consequent release of CQ (**Figure 5D**). Instead, non-illuminated tumours showed no significant changes in the amount of **1C** nor CQ.

## Discussion

CQ and HCQ have emerged as versatile agents with therapeutic potential beyond their original antimalarial applications. Their ability to modulate lysosomal function and inhibit autophagy has attracted significant interest in oncology, particularly for overcoming therapy resistance.^6–8^ Accumulating evidence indicates that CQ and HCQ can selectively target CSCs – a subpopulation responsible for tumour relapse and metastasis – thereby enhancing the efficacy of conventional chemotherapeutic and radiotherapeutic regimens.^14,15^ Through lysosomal alkalinisation and autophagy inhibition, these agents can disrupt key survival pathways in CSCs and bulk tumour cells.^36,37,39^ Together, these findings highlight the promise of CQ and HCQ as adjuvant anticancer agents, though their precise mechanisms and therapeutic window remain subjects of active investigation.

Despite the therapeutic promise of CQ and HCQ, their long-term use in oncology remains constrained by dose-dependent toxicity. Retinopathy, neuromyopathy, and cardiomyopathy have been documented with chronic administration, underscoring the narrow therapeutic window of these agents.^44–47^ Although HCQ offers improved tolerability compared to CQ and is therefore favoured in most clinical settings, achieving pharmacologically effective concentrations in tumours without systemic adverse effects remains challenging. These limitations highlight the need for strategies that enhance spatial and temporal drug selectivity. Photopharmacology provides an elegant solution by enabling light-controlled activation of bioactive molecules with high precision.^44–47^ Through the incorporation of photoswitchable or photocleavable motifs, drug activity can be confined to the tumour site, minimising systemic exposure. Building upon this concept and given the absence of photopharmacological approaches to modulate autophagy inhibition, we explored the design of coumarin-caged CQ and HCQ derivatives. This approach aimed to achieve localised, light-dependent activation of CQ/HCQ to overcome toxicity barriers while maintaining their capacity to target CSCs and disrupt autophagy in cancer models.

The design of photocaged CQ and HCQ analogues aimed to achieve complete suppression of activity in the dark and efficient recovery of the parent drug under visible light. Given that CQ’s pharmacological effect relies on its basic amine centres, caging these functionalities with a photolabile group was expected to block lysosomal accumulation and autophagy inhibition until photoactivation. Incorporation of the DEACM group provided a visible-light-sensitive photocage compatible with biological applications, while also improving absorption at wavelengths of lower phototoxicity. The synthetic approach enabled selective modification at both the aliphatic and aromatic amines, yielding mono- and bis-protected derivatives for both CQ and HCQ.

Photochemical studies confirmed that all DEACM-protected derivatives were stable under dark conditions yet efficiently released their parent drugs upon illumination with visible light. Among them, the aliphatic amine-protected species (**1C** and **1H**) showed the fastest and most efficient uncaging, with t₉₀ of ∼3 min and 68% (H)CQ recovery under 420 nm light – consistent with yields reported for other coumarin-based photocages.^60,61^ The slower photolysis of the aromatic and bis-protected derivatives likely reflects their worse leaving group ability. Notably, **1C** retained meaningful uncaging efficiency beyond its absorption maximum: substantial release occurred at 455–470 nm with short exposures, and even at 500 nm we detected ∼10% CQ release in 2 min. This breadth reflects both DEACM’s tailing absorption and the rapid photolysis of the quaternary adduct, enabling very short illuminations and the use of sub-optimal (but more penetrating) absorbing wavelengths when necessary.

In adherent cancer cell lines, **2C**, **3C**, **2H**, and **3H** were not suitable as light-activated prodrugs as they showed activity in the dark similar to the parent molecules. Instead, caging the aliphatic amine in **1C** and **1H** effectively abolished CQ/HCQ cytotoxicity under dark conditions, confirming that lysosomal accumulation and autophagy inhibition were blocked prior to illumination. CQ-derived **1C** showed a larger window than HCQ-derived **1H**. Upon brief irradiation, **1C** regained potent, dose-dependent cytotoxicity across multiple cancer cell lines covering HNSCC, breast, and colon cancer, recapitulating the effects of their parent drugs. This light-dependent restoration of activity, absent in control experiments with illumination alone, directly links the observed cytotoxicity to photorelease of the active compound. Furthermore, LC3-II accumulation following illumination of **1C** mirrored that induced by CQ, supporting that autophagy inhibition remains the principal mechanism of action post-uncaging.

Importantly, the light-controlled cytotoxicity of **1C** extended to CSCs. We used tumourspheres as a model of CSCs and confirmed this to be enriched in stemness markers. Under dark conditions, **1C** was inert and allowed normal sphere formation, whereas illumination fully prevented spheroid growth and viability, matching the effects of free CQ. Moreover, the selected cell lines appeared to exhibit greater sensitivity to both CQ and **1C** when cultured as spheroids compared to adherent conditions, potentially underscoring an increased susceptibility of CSCs to autophagy inhibition. Collectively, these findings establish **1C** as a potent photosensitive prodrug, enabling precise spatial and temporal control over autophagy inhibition and CSC fate *in vitro*.

*Ex vivo* and *in vivo* optical experiments demonstrated that visible light penetrates tumour tissue sufficiently to trigger uncaging of **1C**. Firstly, we confirmed that UV and blue light (405, 430, and 470 nm) can cross tumour samples of up to 9 mm in thickness *ex vivo*, with blue light (470 nm) exhibiting the highest irradiance. This light effectively induced CQ release in living mice bearing orthotopic breast tumours. A single 10-minute illumination following intratumoural administration of **1C** led to approximately 50% consumption of the photocage and measurable generation of CQ within the tumour, while non-illuminated controls remained unchanged. These results confirm that photochemical activation can occur efficiently in a physiological context under light conditions compatible with biological safety. Previous studies had modelled and studied the optical properties of tumours and malignant tissue.^67–70^ Our work complements these by confirming that, even with scattering and absorption caused by the tumour tissue, clinically relevant light doses can achieve localised activation, supporting the development of photoresponsive therapeutics.

Collectively, these data demonstrate proof of concept for spatially confined activation of a prototypical autophagy inhibitor *in vitro* and *in vivo*. The ability to trigger CQ release within tumour tissue using minimally invasive light exposure highlights the translational potential of this approach. While external illumination will find utility for easily accessible cancers such as breast cancer, HNSCC, and skin cancer, ongoing work with implantable and wireless LEDs and fibres^71–73^ will expand the use to other tissues. Future studies will focus on using other more selective autophagy inhibitors, using red-shifted photocaging groups, and establishing a proof of concept in efficacy studies *in vivo*.

## Conclusions

This study demonstrates a successful strategy to impart light-responsiveness to CQ/HCQ-based autophagy inhibitors through photocaging of their basic amine centres. The use of the DEACM group allowed efficient and reversible control of CQ/HCQ activity under UV and blue light, achieving full suppression of cytotoxicity in the dark and rapid restoration upon illumination. Among the prepared derivatives, the aliphatic amine–caged analogue **1C** emerged as the most effective photoresponsive compound, displaying fast uncaging kinetics and robust light-dependent biological activity.

Photochemical and biological analyses confirmed that **1C** retains the mechanistic hallmark of CQ – autophagy inhibition – while enabling precise spatial and temporal activation. Its ability to eradicate CSC-enriched tumourspheres under illumination underscores the therapeutic potential of light-controlled autophagy modulation to overcome resistance and recurrence. Moreover, *in vivo* activation of **1C** within tumour tissue validates the feasibility of visible-light-triggered drug release under physiologically relevant conditions.

Overall, these results establish the conceptual and practical foundation for photopharmacological control of autophagy inhibitors. The design principles demonstrated here could be extended to other weak-base chemotypes or photoswitchable scaffolds, paving the way toward safer and more selective autophagy-targeted therapies in oncology. Further development towards clinical use could represent a more selective and safer therapy for cancer treatment.

## Acknowledgements

L.J.-C. has received funding from the European Union’s Horizon 2020 research and innovation programme under Marie Sklodowska-Curie Grant Agreement 841089 and from grants IJC2020-043482-I and RyC2023-042477-I funded by MICIU/AEI/10.13039/501100011033 and the European Union NextGenerationEU/PRTR. S.A.M. received funding from a fellowship of the CSIC JAE Intro ICU 2024 program (JAEICU_24_00882). M.E.LL. has received funding from FIS project PI24/00630. The UPLC analysis from the *in vivo* studies were done by Innopharma platform from University of Santiago de Compostela. The authors thank Gemma Fabrias, Mireia Casasampere, and Tania Roda from the Research Unit on BioActive Molecules for support with the Western Blot studies, Yolanda Pérez from the NMR service for analytical and instrumental support, and Elisabet Perez Albadalejo, Jorge Gandia, and Xavier Rovira for biological support.

## Author contributions

Conceptualisation, A.L. and L.J.-C.; data curation, S.A.M, C.S., L.M., M.B.F., B.G.P., S.M.Z., Z.V.D.R., and L.J.-C.; funding acquisition, A.L. and L.J.-C.; methodology, S.A.M, C.S., L.M., M.B.F., B.G.P., S.M.Z., Z.V.D.R., and L.J.-C.; supervision, Y.G.-M., M.E.L.P., Z.V.D.R., A.L., and L.J.-C.; writing – original draft, L.J.-C.; writing – reviewing & editing, all authors.

