## Supplementary information: methods and supplementary figures for "Photocaged chloroquine derivatives for the light-dependent inhibition of autophagy in cancer stem cells"

[1]MCS, Laboratory of Medicinal Chemistry & Synthesis, Department of Biological Chemistry, Institute for Advanced Chemistry of Catalonia (IQAC-CSIC), Jordi Girona 18-26, 08034, Barcelona, Spain.
[2]Synthesis of High Added Value Molecules (SIMChem), Institute for Advanced Chemistry of Catalonia (IQAC-CSIC), Jordi Girona 18-26, 08034, Barcelona, Spain.
[3]Head and neck cancer: Biomedical Research in Tumor Stem Cells Group, Vall d'Hebron Research Institute (VHIR), 08035 Barcelona, Spain.
[4]Clinical Biochemistry, Drug Delivery & Therapy Group (CB-DDT), Vall d'Hebron Institute of Research (VHIR), Vall d'Hebron University Hospital, Vall d'Hebron Barcelona Hospital Campus, Passeig de la Vall d'Hebron, 119-129, 08035 Barcelona, Spain.
[5]Networking Research Center on Bioengineering, Biomaterials and Nanomedicine (CIBER-BBN), Instituto de Salud Carlos III, Barcelona, Spain.
[6]Functional Validation & Preclinical Research (FVPR), Unit20 ICTS Nanbiosis, Vall d'Hebron Institute of Research (VHIR), 08035 Barcelona, Spain.
[7]Drug Discovery & Medicinal Chemistry group, Department of Biological Chemistry, Institute for Advanced Chemistry of Catalonia (IQAC-CSIC), Jordi Girona 18-26, 08034, Barcelona, Spain

### Chemistry methods

#### General chemistry methods

All reagents were obtained from commercial sources and used without further purification. Reactions were conducted under an inert atmosphere of nitrogen or argon, unless not using anhydrous solvents. Anhydrous solvents were obtained from a solvent purification system (PureSolv-EN) and kept under a nitrogen atmosphere.

Analytical thin layer chromatography (TLC) was carried out on aluminum sheets coated with silica gel (Macherey-Nagel, 60F, 0.2 mm, ALUGRAM Sil G/UV254). The spots were visualised by UV irradiation (254 nm) and by staining with a $KMnO_4$ solution followed by heating.

Flash column chromatography was performed on 60 silica gel (Panreac, 40-63 μm particle size) or with a Biotage Isolera One automated system with Biotage KP-C18-SH or Sfär Silica D cartridges. The eluents are specified in each case.

NMR spectra were recorded on a Varian-Mercury 400 MHz spectrometer. Data are reported as follows: chemical shift, multiplicity (s = singlet, d = doublet, t = triplet, q = quartet, m = multiplet), coupling constant, and integration. Chemical shifts (δ) are reported in parts per million (ppm) downfield from TMS and are referenced to the residual solvent peak. Coupling constants (*J*) are quoted in Hertz (Hz).

Low resolution mass spectra (*m/z*) were recorded on a Waters 2795 Alliance coupled to a Diode array detector (Agilent 1100) and an electrospray ionisation (ESI) Quattro Micro, or on a Waters Acquity ARC coupled to an Acquity SQD2 MS detector. Selected peaks are reported in Daltons and their intensities given as percentages of the base peak. High-resolution mass spectra (HRMS) were acquired using a flow injection analysis (FIA) setup with ultrahigh-performance liquid chromatography (UPLC) (Acquity Premier, Waters) coupled to a Select Series Cyclic IMS Q-TOF mass spectrometer (Waters) equipped with a thermostated electrospray ionization (ESI) source. The UPLC system included a binary solvent manager (BSM), sample manager with flow-through needle (SM-FTN), column oven (CM), and photodiode array detector (PDA), all from the Acquity Premier line (Waters). Data from mass spectra were analysed by electrospray ionisation in positive and negative modes using MassLynx 4.1 software (Waters).

Analytical high-performance liquid chromatography (HPLC) was performed on a Thermo Ultimate 3000SD (Thermo Scientific Dionex) coupled to a PDA detector and an LTQ XL ESI-ion trap mass spectrometer (Thermo Scientific) with a Sunfire C18 2.5um 4.6x50mm (Waters) column, or on an ESI Quattro Micro MS detector (Waters) with a ZORBAX Extend-C18 3.5 µm, 2.1 × 50 mm (Agilent) column. The purity of the final compounds was determined to be >95%.

### Synthetic routes and compound characterisation

#### *N*⁴-(7-Chloroquinolin-4-yl)-*N*¹,*N*¹-diethylpentane-1,4-diamine (chloroquine, 5C)

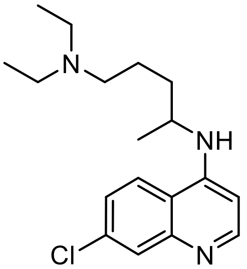

Chloroquine diphosphate (1.00 g, 2.00 mmol) was dissolved in water (6.0 mL), and sodium hydroxide (12% Wt, 3.0 mL, 9.0 mmol) was added. The mixture was stirred at room temperature for 30 min, and then 2 mL of EtOAc were added. After another 30 min, the mixture was extracted with 3xEtOAc, dried over anhydrous $Na_2SO_4$, filtered, and concentrated under reduced pressure to give neutral chloroquine **5C** (656 mg, 2.25 mmol, quant.) as a white solid. $^1$H NMR (400 MHz, DMSO-$d_6$) δ 8.37 (s, 1H), 8.35 (d, *J* = 3.7 Hz, 1H), 7.76 (d, *J* = 2.2 Hz, 1H), 7.42 (dd, *J* = 9.0, 2.3 Hz, 1H), 6.90 (d, *J* = 8.1 Hz, 1H), 6.51 (d, *J* = 5.6 Hz, 1H), 3.80 – 3.64 (m, 1H), 2.40 (q, *J* = 7.1 Hz, 4H), 2.35 (t, *J* = 7.0 Hz, 2H), 1.75 – 1.63 (m, 1H), 1.59 – 1.40 (m, 3H), 1.23 (d, *J* = 6.4 Hz, 3H), 0.90 (t, *J* = 7.1 Hz, 6H). The NMR peaks were in agreement with reported values.[1]

#### (7-(Diethylamino)-2-oxo-2*H*-chromen-4-yl)methyl methanesulfonate (7)

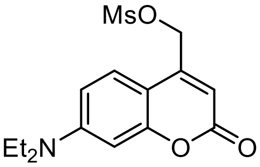

To a solution of 7-(diethylamino)-4-(hydroxymethyl)-2H-chromen-2-one (500 mg, 2.02 mmol) in anhydrous DCM (14.4 mL) were added triethylamine (310 µL, 2.22 mmol) and methanesulfonyl chloride (172 µL, 2.22 mmol). The mixture was stirred at room temperature for 2 h, then diluted with DCM, washed with 1x saturated $NH_4Cl$, 1xsaturated $NaHCO_3$, and 1xbrine, dried over anhydrous $Na_2SO_4$, filtered, and concentrated under reduced pressure to give the mesyl product **7** (655 mg, 2.01 mmol, quant.) as a green solid, which is taken directly to the following step without further purification. $^1$H NMR (400 MHz, $CDCl_3$) δ 7.34 (d, *J* = 9.0 Hz, 1H), 6.66 (d, *J* = 9.0 Hz, 1H), 6.57 (d, *J* = 2.6 Hz, 1H), 6.19 (s, 1H), 3.43 (q, *J* = 7.1 Hz, 4H), 3.11 (s, 3H), 1.22 (t, *J* = 7.1 Hz, 7H); *m/z* (ESI+) 326.2 ($MH^+$, 100%). The NMR peaks were in agreement with reported values.[2]

#### 4-(Bromomethyl)-7-(diethylamino)-2*H*-chromen-2-one (8)

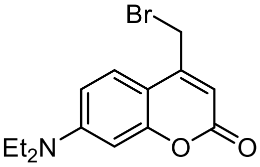

Lithium bromide (524 mg, 6.04 mmol) was added to a solution of mesylate **7** (655 mg, 2.01 mmol) in anhydrous THF (28.7 mL). The mixture was stirred at room temperature overnight, and then concentrated under reduced pressure. The residue was redissolved in DCM, washed with 1xwater, dried over anhydrous $Na_2SO_4$, filtered, and concentrated under reduced pressure to give bromide **8** (624 mg, 2.01 mmol, quant.) as a green oil. $^1H$ NMR (400 MHz, $CDCl_3$) δ 7.50 (d, *J* = 9.0 Hz, 1H), 6.63 (d, *J* = 7.3 Hz, 1H), 6.52 (s, 1H), 6.14 (s, 1H), 4.40 (s, 2H), 3.42 (q, *J* = 7.1 Hz, 4H), 1.22 (t, *J* = 7.1 Hz, 6H); *m/z* (ESI+) 312.1 (MH+, 100%). The NMR peaks were in agreement with reported values.[3]

#### 4-(((7-Chloroquinolin-4-yl)(5-(diethylamino)pentan-2-yl)amino)methyl)-7-(diethylamino)-2*H*-chromen-2-one formate (1C), 4-((7-chloroquinolin-4-yl)amino)-*N*-((7-(diethylamino)-2-oxo-2*H*-chromen-4-yl)methyl)-*N,N*-diethylpentan-1-aminium formate (2C), and 4-((7-chloroquinolin-4-yl)((7-(diethylamino)-2-oxo-2*H*-chromen-4-yl)methyl)amino)-*N*-((7-(diethylamino)-2-oxo-2*H*-chromen-4-yl)methyl)-*N,N*-diethylpentan-1-aminium formate (3C)

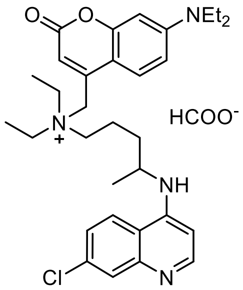

**1C**

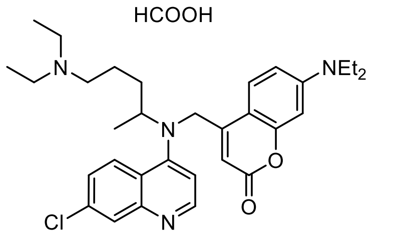

**2C**

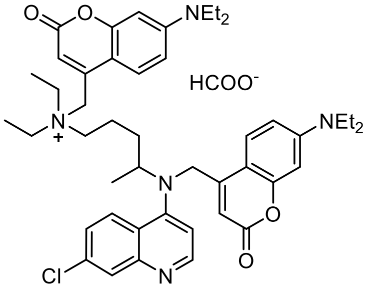

**3C**

To neutral chloroquine **5C** (116 mg, 0.363 mmol) was added a solution of bromide **8** (124 mg, 0.400 mmol) in anhydrous acetonitrile (3.6 mL). The mixture was stirred at 60 °C overnight. The mixture was concentrated under reduced pressure. The crude product was purified by C18 flash column chromatography (10% to 100% MeCN with 0.05% FA in water with 0.05% FA) to give **1C** (22 mg, 0.038 mmol, 10%), **2C** (26 mg, 0.044 mmol, 12%), and **3C** (9.4 mg, 0.011 mmol, 3%).

**1C**: $^1H$ NMR (400 MHz, DMSO-$d_6$) δ 8.52 (t, *J* = 7.5 Hz, 2H), 8.13 (s, 0.4H, *H*COOH), 7.89 (d, *J* = 2.2 Hz, 1H), 7.80 (d, *J* = 9.2 Hz, 1H), 7.69 (d, *J* = 9.0 Hz, 1H), 6.76 (s, 1H), 6.69 (dd, *J* = 9.2, 2.6 Hz, 1H), 6.56 (d, *J* = 2.6 Hz, 1H), 6.24 (s, 1H), 4.64 – 4.52 (m, 2H), 3.94 (s, 1H), 3.55 – 3.33 (m, 10H), 1.84 – 1.72 (m, 2H), 1.72 – 1.57 (m, 1H), 1.57 – 1.43 (m, 1H), 1.32 – 1.22 (m, 9H), 1.10 (t, *J* = 7.0 Hz, 6H); $^{13}C$ NMR (101 MHz, MeOD) δ 162.4, 157.9, 156.8, 152.9, 144.3, 144.1, 140.8, 140.5, 128.5, 127.1, 126.5, 120.5, 117.1, 114.8, 110.9, 108.8, 100.1, 98.7, 60.1, 57.8, 56.5, 51.0, 45.7, 33.2, 20.8, 19.9, 12.7, 8.7; *m/z* (ESI+) 549.4 (M+, 25%); HRMS (*m/z*): [M]+ calcd. for C32H42N4O2Cl+, 549.2991, found 549.2994.

**2C**: $^{1}$H NMR (400 MHz, MeOD) δ 8.74 (d, *J* = 9.1 Hz, 1H), 8.59 (d, *J* = 7.2 Hz, 1H), 8.35 (s, 1H, *H*COOH), 7.89 (d, *J* = 1.7 Hz, 1H), 7.77 (dd, *J* = 9.0, 1.5 Hz, 1H), 7.71 (d, *J* = 9.1 Hz, 1H), 7.14 (d, *J* = 7.5 Hz, 1H), 6.86 (dd, *J* = 9.1, 2.5 Hz, 1H), 6.59 (d, *J* = 2.5 Hz, 1H), 6.06 (s, 1H), 5.10 (s, 1H), 4.27 (q, *J* = 6.4 Hz, 1H), 3.52 (q, *J* = 7.1 Hz, 4H), 3.28 – 3.15 (m, 6H), 2.08 – 1.80 (m, 4H), 1.48 (d, *J* = 6.4 Hz, 3H), 1.32 (t, *J* = 7.2 Hz, 6H), 1.24 (t, *J* = 7.0 Hz, 6H); $^{13}$C NMR (101 MHz, MeOD) δ 167.8, 163.4, 157.6, 157.3, 153.0, 151.5, 149.1, 142.3, 140.6, 128.9, 127.6, 126.2, 118.8, 118.1, 110.7, 106.7, 105.0, 101.1, 98.3, 54.7, 52.8, 51.7, 48.3, 45.7, 33.6, 22.1, 20.0, 12.7, 9.1; *m/z* (ESI+) 549.4 (MH+, 18%); HRMS (*m/z*): [MH]+ calcd. for C32H42N4O2Cl+, 549.2991, found 549.2993.

**3C**: $^{1}$H NMR (400 MHz, MeOD) δ 8.73 (s, 1H), 8.58 (s, 1H), 8.29 (s, 2H), 7.88 (s, 1H), 7.82 – 7.64 (m, 3H), 6.86 (d, *J* = 8.6 Hz, 1H), 6.80 (s, 1H), 6.59 (d, *J* = 2.3 Hz, 1H), 6.50 (d, *J* = 2.0 Hz, 1H), 6.20 (s, 1H), 6.12 – 6.00 (m, 1H), 5.11 (s, 1H), 3.57 – 3.42 (m, 10H), 1.49 – 1.37 (m, 4H), 1.31 – 1.14 (m, 15H); $^{13}$C NMR (101 MHz, MeOD) δ 165.6, 162.0, 161.1, 157.2, 156.5, 156.2, 155.8, 151.7, 151.6, 147.8, 142.8, 140.9, 127.6, 126.4, 125.7, 124.8, 123.0, 117.4, 116.7, 113.4, 109.5, 109.3, 107.4, 105.3, 103.8, 99.7, 97.4, 97.0, 66.3, 58.7, 56.4, 55.1, 48.5, 44.3, 44.3, 31.8, 29.4, 19.4, 18.5, 11.3, 11.3, 7.3; *m/z* (ESI+) 778.4 (M+, 30%); HRMS (*m/z*): [M]+ calcd. for C46H57N5O4Cl+, 778.4094, found 778.4113.

##### $N^{4}$-(7-Chloroquinolin-4-yl)-$N^{1}$,$N^{1}$-diethylpentane-1,4-diamine (hydroxychloroquine, 5H)

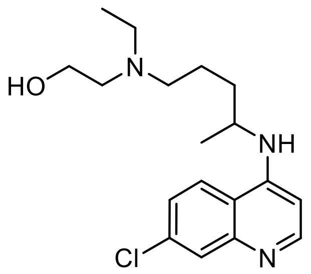

To a suspension of hydroxychloroquine sulphate (420 mg, 0.968 mmol) in chloroform (8.4 mL) was added 30% ammonium hydroxide until basic pH, which gave a clear solution that was stirred for another 1 h. The organic layer was washed with 2xwater, dried over anhydrous $Na_2SO_4$, filtered, and concentrated under reduced pressure to give neutral hydroxychloroquine **5H** (330 mg, 0.968 mmol, quant.) as a colourless oil. $^{1}$H NMR (400 MHz, $CDCl_3$) δ 8.51 (d, *J* = 5.4 Hz, 1H), 7.95 (d, *J* = 1.9 Hz, 1H), 7.85 – 7.69 (m, 1H), 7.37 (dd, *J* = 8.9, 1.9 Hz, 1H), 6.40 (d, *J* = 5.5 Hz, 1H), 5.21 – 4.94 (m, 1H), 3.78 – 3.69 (m, 1H), 3.66 – 3.53 (m, 2H), 2.77 – 2.50 (m, 6H), 1.72 – 1.60 (m, 4H), 1.33 (d, *J* = 6.4 Hz, 3H), 1.07 (q, *J* = 8.1 Hz, 3H); *m/z* (ESI+) 336.4 (MH+, 30%). The NMR peaks were in agreement with reported values.[4]

**4-(((7-Chloroquinolin-4-yl)(5-(ethyl(2-hydroxyethyl)amino)pentan-2-yl)amino)methyl)-7-(diethylamino)-2*H*-chromen-2-one (2*H*), and 4-((7-chloroquinolin-4-yl)((7-(diethylamino)-2-oxo-2*H*-chromen-4-yl)methyl)amino)-*N*-((7-(diethylamino)-2-oxo-2*H*-chromen-4-yl)methyl)-*N*-ethyl-*N*-(2-hydroxyethyl)pentan-1-aminium (3H)**

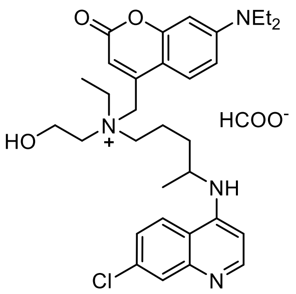

**1H**

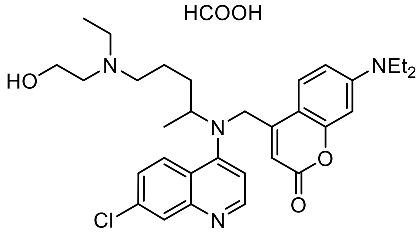

**2H**

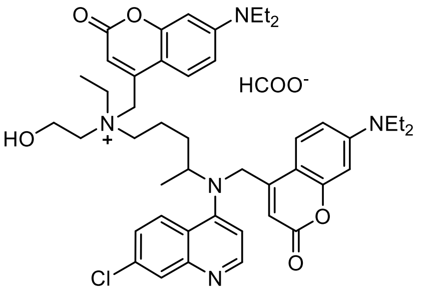

**3H**

To neutral hydroxychloroquine **5H** (110 mg, 0.328 mmol) was added a solution of bromide **8** (110 mg, 0.328 mmol) in anhydrous acetonitrile (3.3 mL). The mixture was heated to 60 °C overnight and then concentrated under reduced pressure. The crude product was purified by C18 flash column chromatography (10% to 100% MeCN with 0.05% FA in water with 0.05% FA) to give **1H** (58 mg, 0.0376 mmol, 18%), **2H** (68 mg, 0.12 mmol, 37%), and **3C** (60 mg, 0.075 mmol, 23%).

**1H**: $^{1}$H NMR (400 MHz, MeOD) δ 8.49 (dd, *J* = 9.1, 2.3 Hz, 1H), 8.45–8.38 (m, 1H), 7.90 (s, 1H), 7.77 (d, *J* = 9.2 Hz, 1H), 7.66 (dd, *J* = 9.0, 1.5 Hz, 1H), 6.87 (d, *J* = 6.5 Hz, 1H), 6.80 (dt, *J* = 9.1, 3.0 Hz, 1H), 6.53 (dd, *J* = 4.4, 2.6 Hz, 1H), 6.32 (d, *J* = 8.1 Hz, 1H), 4.86 (s, 2H), 4.15–4.03 (m, 3H), 3.70–3.54 (m, 4H), 3.53–3.44 (m, 6H), 2.08–1.91 (m, 2H), 1.87–1.76 (m, 1H), 1.76–1.65 (m, 1H), 1.45 (t, *J* = 7.0 Hz, 3H), 1.41 (d, *J* = 6.1 Hz, 3H), 1.25–1.18 (m, 6H); $^{13}$C NMR (101 MHz, MeOD) δ 162.5, 162.5, 157.9, 156.3, 152.9, 145.2, 144.2, 141.6, 140.3, 128.3, 127.1, 126.1, 121.4, 117.0, 115.1, 110.8, 108.9, 100.0, 98.8, 62.0, 61.0, 59.0, 57.6, 56.9, 50.8, 45.7, 33.2, 20.9, 19.9, 12.7, 8.8; *m/z* (ESI+) 565.5 (M+, 33%); HRMS (*m/z*): [MH]+ calcd. for C32H42N4O3Cl+, 565.2940, found 565.2922.

**2H**: $^{1}$H NMR (400 MHz, MeOD) δ 8.79 (d, *J* = 9.1 Hz, 1H), 8.62 (d, *J* = 7.6 Hz, 1H), 7.87 (d, *J* = 1.8 Hz, 1H), 7.78–7.68 (m, 2H), 7.17 (d, *J* = 7.8 Hz, 1H), 6.85 (dd, *J* = 9.1, 2.5 Hz, 1H), 6.56 (d, *J* = 2.5 Hz, 1H), 6.16–6.01 (m, 2H), 5.10 (s, 1H), 4.31 (q, *J* = 6.5 Hz, 1H), 3.91–3.83 (m, 2H), 3.51 (q, *J* = 7.0 Hz, 4H), 3.37–3.24 (m, 6H), 2.09–1.78 (m, 4H), 1.49 (d, *J* = 6.4 Hz, 3H), 1.34 (t, *J* = 7.3 Hz, 3H), 1.23 (t, *J* = 7.0 Hz, 6H); $^{13}$C NMR (101 MHz, MeOD) δ 163.4, 157.5, 157.2, 153.0, 151.5, 149.2, 142.2, 140.6, 128.8, 127.7, 126.3, 118.8, 118.0, 110.7, 106.6, 105.0, 101.2, 98.2, 56.7, 55.3, 54.8, 53.7, 51.7, 49.8, 45.7, 33.5, 21.8, 20.1, 12.8, 9.2; *m/z* (ESI+) 565.5 (MH+, 100%); HRMS (*m/z*): [MH]+ calcd. for C32H42N4O3Cl+, 565.2940, found 565.2943.

**3H**: $^{1}$H NMR (400 MHz, MeOD) δ 8.72 (dd, *J* = 9.0, 3.4 Hz, 1H), 8.61 (d, *J* = 7.2 Hz, 1H), 8.41 (s, 1H), 7.90 (s, 1H), 7.81 (dd, *J* = 9.2, 3.4 Hz, 1H), 7.76–7.64 (m, 2H), 7.13 (d, *J* = 7.1 Hz, 1H), 6.86 (dd, *J* = 9.1, 2.4 Hz, 1H), 6.81 (dt, *J* = 9.1, 3.0 Hz, 1H), 6.58 (d, *J* = 2.4 Hz, 1H), 6.48 (dd, *J* = 4.0, 2.6 Hz, 1H), 6.32 (d, *J* = 10.9 Hz, 1H), 6.16–6.01 (m, 2H), 5.13 (s, 1H), 4.28 (q, *J* = 6.9 Hz, 1H), 4.20–4.00 (m, 2H), 3.73–3.40 (m, 12H), 2.14–1.98 (m, 2H), 1.97–1.85 (m, 1H), 1.82–1.71 (m, 1H), 1.53–1.39 (m, 6H), 1.28–1.10 (m, 16H); $^{13}$C NMR (101 MHz, MeOD) δ 168.2, 163.3, 162.5, 162.5, 157.9, 157.6, 157.2, 153.0, 152.9, 151.4, 149.2, 144.2, 142.2, 140.6, 128.9,

127.6, 127.2, 126.3, 118.8, 118.1, 115.1, 110.9, 110.7, 108.9, 106.7, 105.1, 101.2, 98.7, 98.3, 62.0, 61.0, 59.1, 57.7, 57.0, 54.8, 51.6, 45.7, 33.3, 21.0, 20.0, 12.8, 9.0; *m/z* (ESI+) 794.6 (MH+, 45%); HRMS (*m/z*): [MH]+ calcd. for C46H57N5O5Cl+, 794.4043, found 794.4042.

### Photochemistry methods

#### General photochemistry details

96-well LED array plates (LEDA Teleopto) of 405 and 420 nm were used to illuminate the samples for uncaging. For uncaging studies by HPLC or absorbance, the maximum power of 19 mW/cm$^2$ (for 405 nm) and 13 mW/cm$^2$ (for 420 nm) was used. Potencies were measured using a Thorlabs PM100D power energy meter connected to a standard photodiode power sensor (S120VC).

#### Uncaging monitoring by HPLC

UV–vis absorption spectra of a 200 µL solution of each compound at 100 µM in PBS (0.5% DMSO) were recorded using an Infinite M1000 Tecan microplate reader (λ = 300–800 nm).

200 µL solutions of each compound at 100 µM in DMSO or PBS were illuminated for different times, and the corresponding proportion of cage and CQ/HCQ was then measured with a Thermo Ultimate 3000SD instrument (Thermo Scientific Dionex) coupled to a PDA detector and an LTQ XL ESI-ion trap mass spectrometer (Thermo Scientific). Previous calibration curves with each compound were done with at least 4 concentrations, curves were fit by plotting the peak area of the analyte versus the concentrations of the analyte with least-squares linear regression. Uncaging rates are plotted as concentration of cage and CQ/HCQ over the irradiation time.

#### Uncaging monitoring by absorbance

200 µL solutions of each compound at 100 µM in PBS in transparent flat-bottom 96-well plates were illuminated for different times, and the resulting absorbance spectra were measured between 300 nm and 800 nm with 2 nm fixed intervals using a Tecan microplate reader.

### Biology methods

#### Cell lines and culture conditions

The male pharyngeal cancer cell line HTB43 (FaDu) and the female pharyngeal cancer cell line CCL138 were purchased from the American Type Culture Collection (ATCC, Manassas, VA, USA) and were maintained in adherent culture in MEM supplemented with 10% fetal bovine serum (FBS), 1 mM sodium pyruvate, and 1% penicillin-streptomycin (Pen-Strep, final 100 U/mL).

The male laryngeal cancer cell line JHU029 Lucires-GFP was kindly provided by Dr S. Aznar-Benitah (Institute for Research in Biomedicine Barcelona, Spain) and was maintained in adherent culture in RPMI 1640 supplemented with 10% fetal bovine serum (FBS), 1% L-glutamine, and 1% Pen-Strep.

The female breast cancer cell lines MCF7 and MDA-MB-468 were purchased from ATCC. Both cell lines were maintained in adherent culture in DMEM supplemented with 10% FBS.

The female breast cancer cell line MDA-MB-231 was purchased from ATCC and maintained in adherent culture in DMEM-F12 supplemented with 10% FBS, 1 mM sodium pyruvate, and 1% Pen-Strep.

The female colorectal cancer cell line HT29 was purchased from ATCC and maintained in adherent culture in DMEM supplemented with 10% FBS.

#### Cell viability assays in adherent culture

Stock solutions of the photocages were prepared at 40 mM in DMSO and stored at -20 °C. Stock solutions of CQ and HCQ were prepared at 20 mM in water and stored at -20 °C.

Cells were seeded in 96-well tissue culture plates (Nucleon Delta Surface, Thermo Scientific) at a density of $1x10^4$ cells/well, in a volume of 95 µL per well. Then, 5 µL of compound solutions (x20 of desired concentration) were added. Serial dilutions were carried out in cell medium prior to use in each experiment.

Illumination of selected wells was done with a 96-well LED array plate (LEDA Teleopto) at 420 nm placed below the seeded plate for 30-40 s at 8 $mW/cm^2$.

Cells were incubated with the compounds in the dark for a further 3 days, keeping the incubator closed to prevent light exposure. Viability was determined with a CellTiter 96 assay (MTS, Promega), according to the manufacturer's instructions. Briefly, 15 µL of CellTiter 96 reagent was added to each well, the plates were incubated at 37 °C for 2 h, and the absorbance at 490 nm was recorded using a Tecan Spark 20M Multimode Microplate reader. Percentage of live cells is expressed with respect to the non-treated cells.

#### Western blot

For LC3-II protein analysis, $5x10^5$ cells were plated in 6-well plates and allowed to adhere for 48 h. Cells were treated for 2-4 h with the photocages or CQ/HCQ. After treatment, cells were detached with trypsin-EDTA and collected in pellets. The pellets were resuspended in an aqueous solution of 0.25 M sucrose and submitted to three cycles of 5 s sonication (probe)/5 s resting on ice. The cell lysate was centrifuged at 2500 g for 5 min. The supernatant was collected, and protein concentration was estimated (BSA as a standard) using a BCA protein determination kit (Thermo Scientific) according to the manufacturer's instructions.

Samples were loaded onto a 12% polyacrylamide gel, separated by electrophoresis at 100 V/1.5-2.5 h and transferred onto a PVDF membrane (100 V/2 h). Unspecific binding sites were then blocked with 5% milk in TBST. Anti-LC3 from Abcam (code ab48394) was diluted 1:1000 in 5% milk in TBST. Anti-β-actin antibody from Sigma (code A2228) was diluted 1:2000 in 5% milk in TBST. Membranes were incubated overnight at 4 ºC under gentle agitation. After washing with TBST, membranes were probed with the corresponding secondary antibody for 1 h at room temperature (LC3: anti-rabbit (NA934V), diluted 1:1000 in 5% milk in TBST; β-actin: anti-mouse (170-6516), diluted 1:10000 in 5% milk in TBST). Antibody excess was eliminated by washing with TBST, and protein detection was carried out using ECL and membrane scanning with LI-COR C-DiGit® Blot Scanner. Band intensities were quantified by LI-COR Image Studio Lite Software.

#### Spheres

To grow CSC-enriched spheres, cells were grown in 96-well ultra-low attachment plates (Corning) in 3D tumorsphere medium XF (PromoCell).

Cells were seeded at a density of 4-8x$10^4$ cells/well, in a volume of 95 µL per well. Then, 5 µL of compound solutions (x20 of desired concentration) were added. Serial dilutions were carried out in cell medium prior to use in each experiment. Illumination of selected wells was done with a 96-well LED array plate (LEDA Teleopto) at 420 nm placed below the seeded plate for 40 s at 8 mW/$cm^2$.

Cells were grown for 7-10 days and pictures were taken with an EVOS microscope M700.

To assess the viability of the spheres, to each well was added 20 µL of CellTiter 96 reagent (MTS assay, Promega). Then, the plates were incubated at 37 °C for 24 h, and the absorbance at 490 nm was recorded using a Tecan Spark 20M Multimode Microplate reader. Percentage of live cells is expressed with respect to the non-treated cells.

To grow spheres of second and third generations, spheres formed were collected in a falcon tube and sedimented by gravity for 15 minutes, discarding individual cells. Once sedimented, the supernatant was removed and spheres were resuspended in TrypLE™ Express trypsin (Gibco, Life Technologies) and left at 37 °C for 10 minutes, mixing after 5 minutes. The solution was diluted with 5x PBS, centrifuged, resuspended in sphere growth medium, and seeded again under the same initial conditions to form the next generation of CSCs.

#### qRT-PCR

Total RNA extraction from JHU029 and JHU029-derived CSCs was performed using Direct-zol RNA Miniprep Kits (Zymo Research) according to the manufacturers' instructions. The cDNA was obtained using the RevertAid H minus first strand cDNA synthesis kit (Thermo Scientific) and a Veriti 96-well thermal cycler (Applied Biosystems). mRNA expression was analyzed using Taqman probes for the following genes: *SOX2* (Hs01053049_s1), *ALDH1A1* (Hs00946916_m1), *CD44* (Hs01075861_m1), *NANOG* (Hs04399610_g1), *OCT3/4* (Hs04260367_gH), and *SNAI1* (Hs00195591_m1), and *IPO8* (Hs00183533_m1) as an endogenous control; in a LightCycler 480 instrument (Roche) using TaqMan™ Universal Master Mix II (Applied Biosystems, CA, USA; Thermo Fisher Scientific). Each sample was run in triplicate and the quantification method used was 2-ΔΔCt.

#### Illumination of tumour samples

Mounted LEDs from Thorlabs (405 nm: M405L4, 430 nm: M430L5, 470 nm: M470L5) with a collimator controlled with a LED Driver (CD2200) were used for illumination of tumours both *ex vivo* and *in vivo* with the setup shown in **Figure S15**. The light crossing the tumour was measured with a photodiode power sensor S120VC connected to a PM100D power energy meter.

Illumination was done at maximum intensity, which corresponded to 84 mW/cm$^2$ for 405 nm, 108 mW/cm$^2$ for 430 nm, and 135 mW/cm$^2$ for 470 nm.

#### Photolysis study *in vivo*

All animal experiments were conduct in accordance with protocols approved by the Ethical Committee for the Use of Experimental Animals (CEEA) at the Vall d'Hebron Research Institue (VHIR), and the local government (CEA-OH/11791/2). *In vivo* studies were carried out by FVPR/U20 of ICTS "NANBIOSIS". Female athymic nude mice (Hsd:Athymic Nude-Foxn1$^{nu}$, Envigo) with orthotopic MDA-MB-231 tumours were used in this study. Briefly, 2.0 × 10$^6$ MDA-MB-231 cells were suspended in a 1:1 mixture of PBS1x:Matrigel and injected into the mammary fat pad (i.m.f.p) in 7 weeks old female mice. Animal welfare, body weight and tumour volume were monitored twice per week. When tumour volumes reached 80–120 mm$^3$, mice were randomised into 3 groups (two animals per group): (1) untreated, (2) treated with **1C**, (3) treated with **1C** and illuminated for 10 minutes. 25 µL of **1C** was administered intratumourally (i.t.) at single dose of 10 mg/kg (in PBS with 5% DMSO). After 10 minutes of treatment, mice were euthanised and tumours were excised, weighted, and homogenised with PBS.

To analyse the samples by HPLC, these were diluted in water containing 0.5% formic acid and then mixed with three volumes of acetonitrile containing the internal standard. The samples were centrifuged, and the supernatant was analysed by UPLC in an ACQUITY UPLC H-Class and Xevo TQD MS System. A HSST3 1.7µm 2.1x100mm (Waters) column was used at a flow rate of 0.6 mL/min, with water + 0.1% formic acid and acetonitrile + 0.1% formic acid as the solvents. Electrospray ionisation was run in positive mode with a source temperature of 150 °C and a desolvation temperature of 600 °C. Capillary voltage was set to 3 kV and the cone voltage was set to 30 V. Desolvation gas flow was 1100 L/h and cone gas flow was set to 150 L/h. 1C and CQ were monitored in multiple reaction monitoring mode.

### Quantification and statistical analysis

All key experiments were independently repeated at least three times with consistent results. All data meeting experimental requirements were included in the analyses, and no data were excluded except in cases of clear technical failure (e.g., sample loss or instrument malfunction). Graph Pad Prism 9 was used to conduct statistical analysis. Data are presented as the mean ± standard deviation (SD).

For the intratumoural photolysis experiment, statistical significance was calculated with a one-way ANOVA with multiple comparisons, and a p value of $p < 0.05$ was considered statistically significant. In all cases, ns: not significant, $p \geq 0.05$; $*p < 0.05$; $**p < 0.01$; $***p < 0.001$; $****p < 0.0001$.

### Supplementary figures

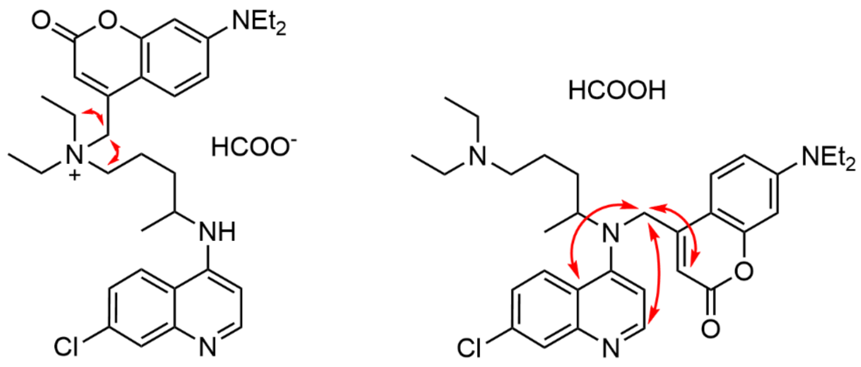

**1C** **2C**

**Figure S1**. Selected HMBC correlations of **1C** and **2C**.

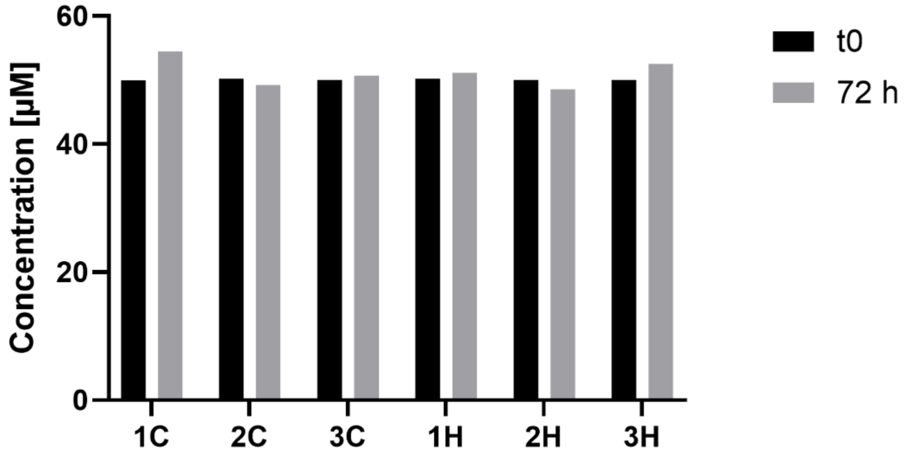

**Figure S2**. Concentration of photocages **1C-3C** and **1H-3H** at 50 µM in DMEM with 10%FBS after 72 h in the dark inside an incubator at 37 °C with 5% DMSO.

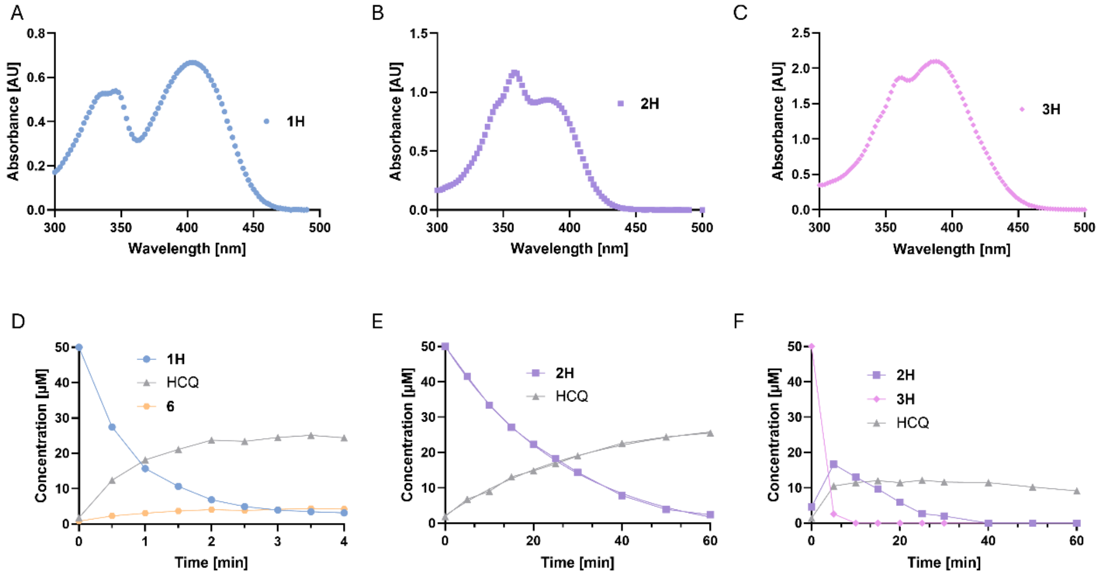

**Figure S3**. Photochemical characterisation of compounds **1H-3H**. (A-C) Absorbance spectra of compounds **1H-3H**, at 100 µM in PBS. (D-F) HPLC quantification of photocage and CQ during illumination at 405 (**2H** and **3H**, 19 mW/cm$^2$) or 420 nm (**1H**, 13 mW/cm$^2$) for the indicated times, initial solution at 50 µM in PBS.

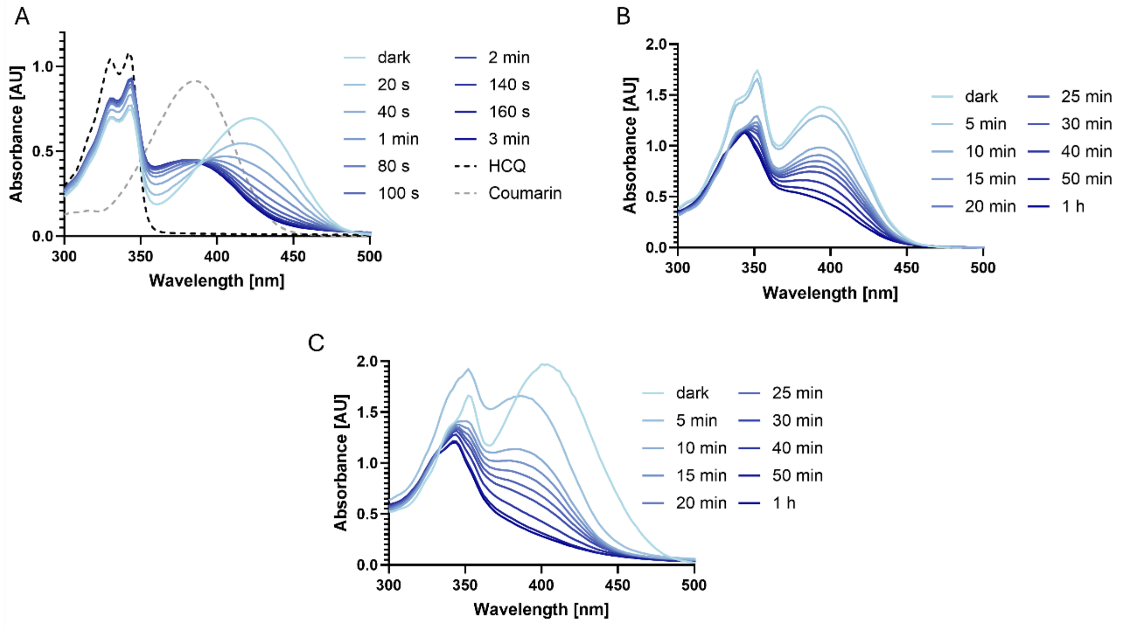

**Figure S4**. A) Changes of the UV-Vis absorption of **1H** recorded in 20-second intervals during 420 nm (13 mW/cm$^2$) irradiation at 100 µM in PBS, dotted lines show the UV-Vis absorption of HCQ and coumarin **6** at 100 µM in PBS. B) Changes of the UV-Vis absorption of **2H** recorded in 5-10-minute intervals during 405 nm (19 mW/cm$^2$) irradiation at 100 µM in PBS. C) Changes of the UV-Vis absorption of **3H** recorded in 5-10-minute intervals during 405 nm (19 mW/cm$^2$) irradiation at 100 µM in PBS.

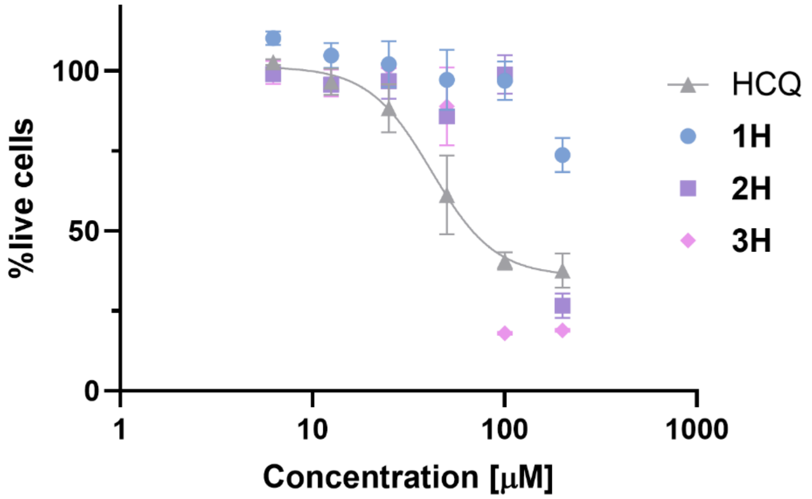

**Figure S5**. Dose-response curves of HCQ and the HCQ photocages **1H-3H** on the viability of MCF7 cells (MTS assay) under dark conditions for 72 h ($IC_{50}$ (CQ) = 41.7 µM; $IC_{50}$ (**3H**) = 55.1 µM). Values are represented as mean ± SD.

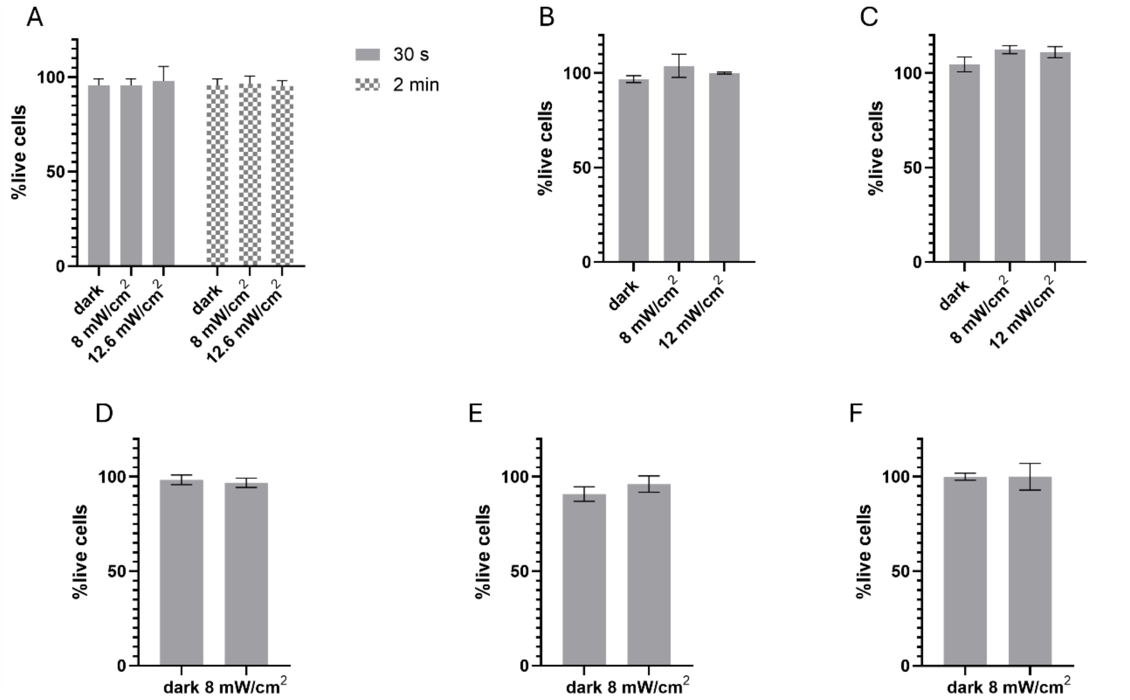

**Figure S6**. Viability of A) MCF7, B) MDA-MB-231, C) MDA-MB-468, D) JHU029, E) HTB43, and F) CCL138 cells after being illuminated for 40 s (and MCF7 also for 2 min) at 420 nm at the indicated potencies, and subsequently incubated for 72 h in the dark. Values are represented as mean ± SD.

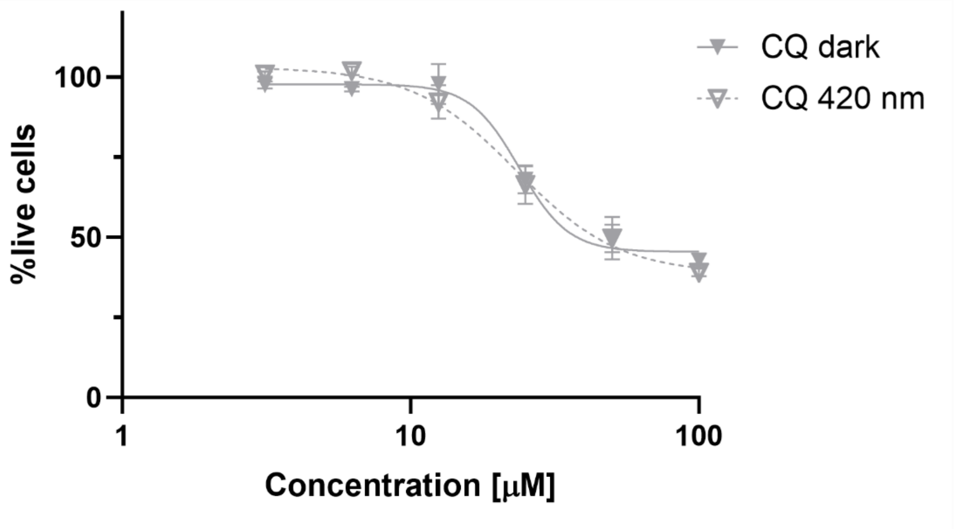

**Figure S7**. Dose-response curves of CQ on the viability of MCF7 cells (MTS assay) under dark vs illumination (420 nm, 8 mW/cm$^2$, 40 s) conditions for 72 h. Values are represented as mean ± SD.

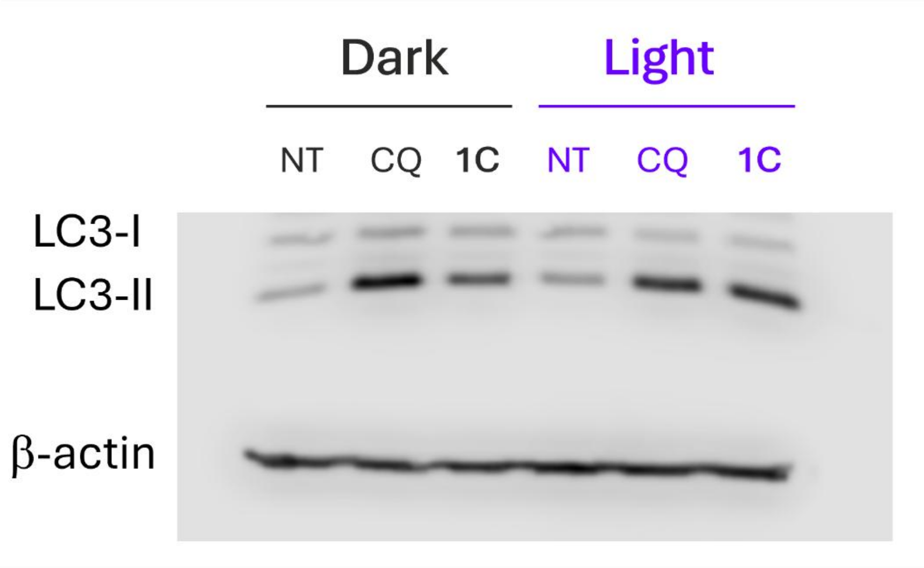

**Figure S8**. Representative western blot of LC3-I and LC3-II after treatment with HCQ or **1C** under dark or illumination conditions (420 nm, 8 mW/cm$^2$, 1 min) in MCF7 cells, actin was used as loading control.

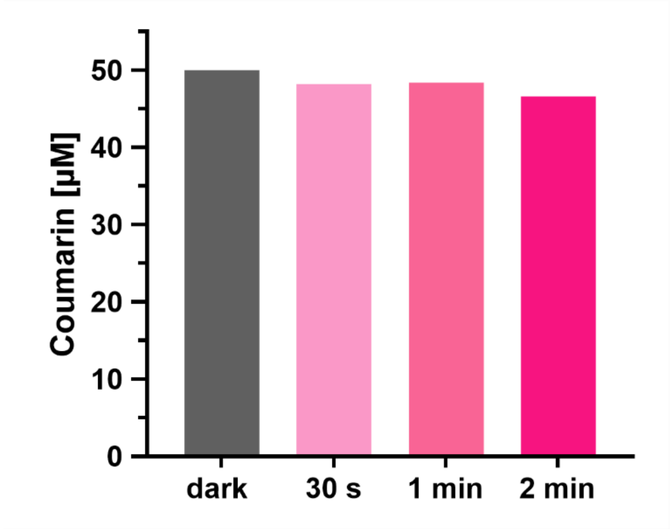

**Figure S9**. Stability of a solution of coumarin alcohol **6** at 50 µM in DMEM after illumination at 420 nm for the indicated times, determined by HPLC.

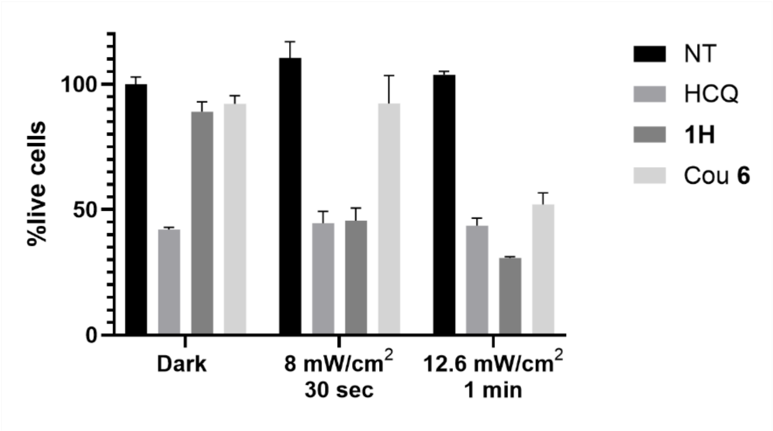

**Figure S10**. Viability of MCF7 cells (MTS assay) treated with HCQ, **1H**, or coumarin **6** at 100 µM for 72 h, under either dark or illumination conditions at the indicated intensity and time. Values are represented as mean ± SD.

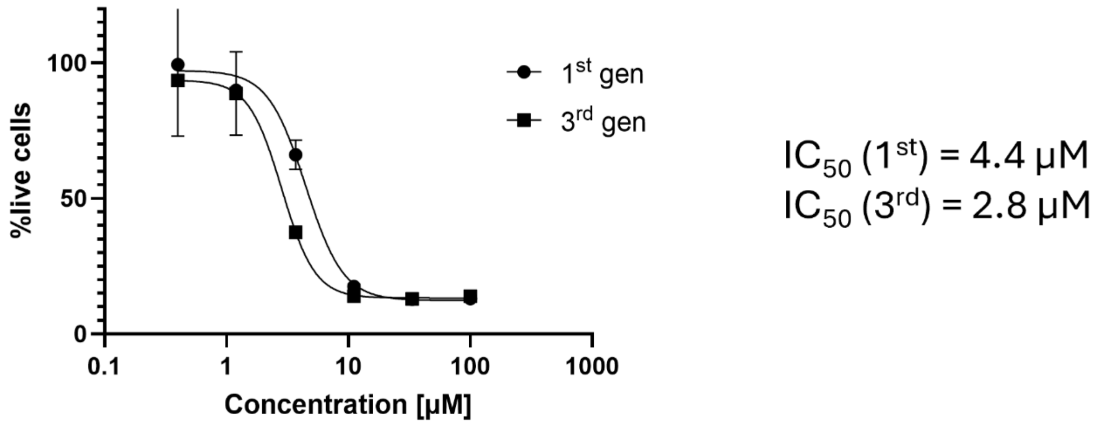

**Figure S11**. Dose-response curves of HCQ on the viability of MCF7 cells (MTS assay) grown as spheres for either 1 or 3 generations for 7-10 days.

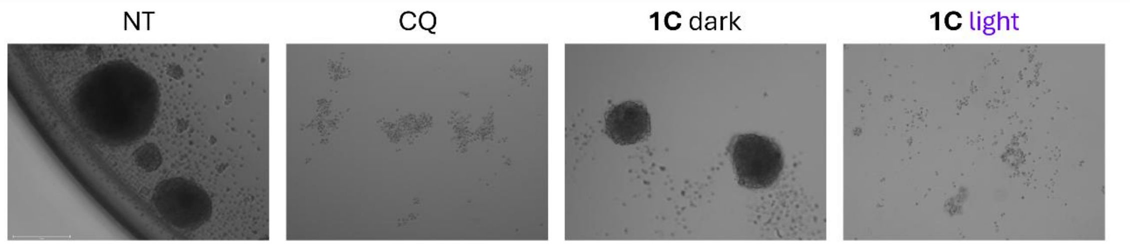

**Figure S12**. Brightfield images of JHU029 cells grown in sphere culture, treated with **1C** at 100 μM under dark vs illumination (420 nm, 8 mW/cm$^2$, 40 s) conditions and of CQ for 10 days.

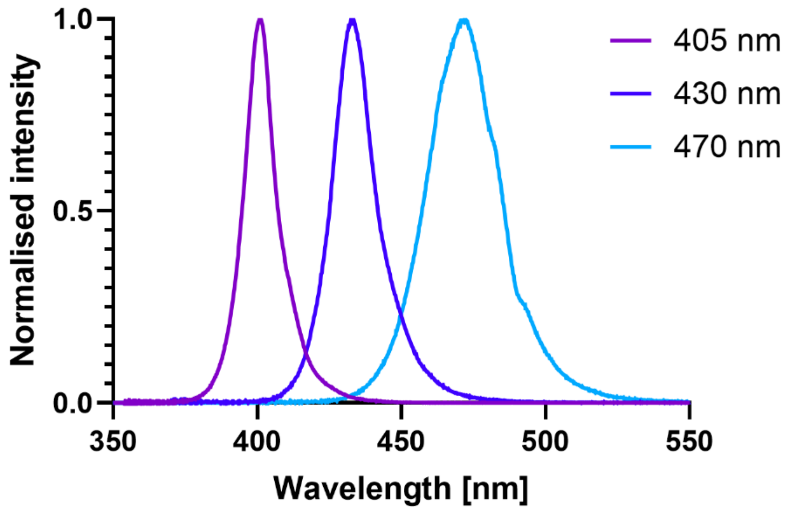

**Figure S13**. Spectral range of tested LEDs.

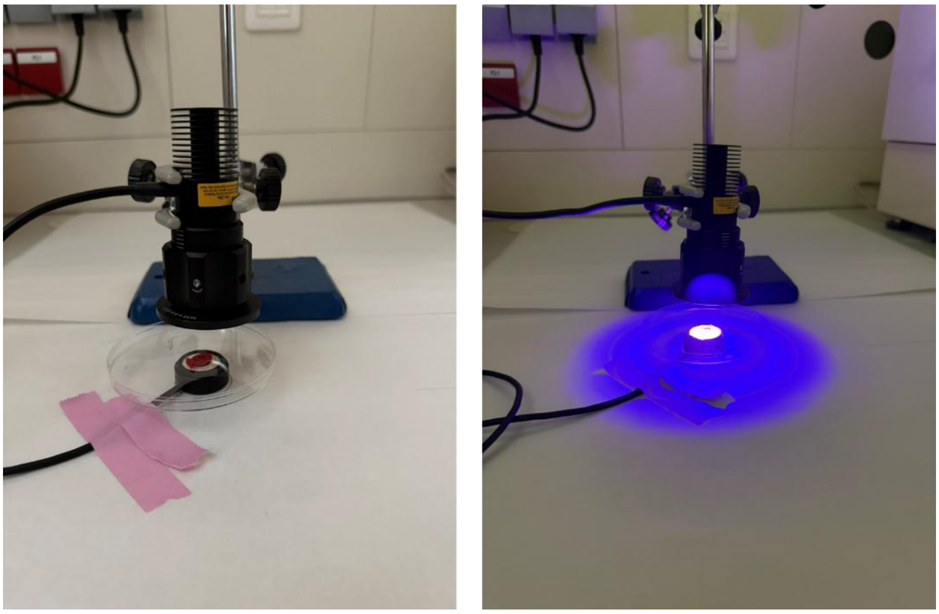

**Figure S14**. Setup for the illumination of tumour samples and irradiance measurement.

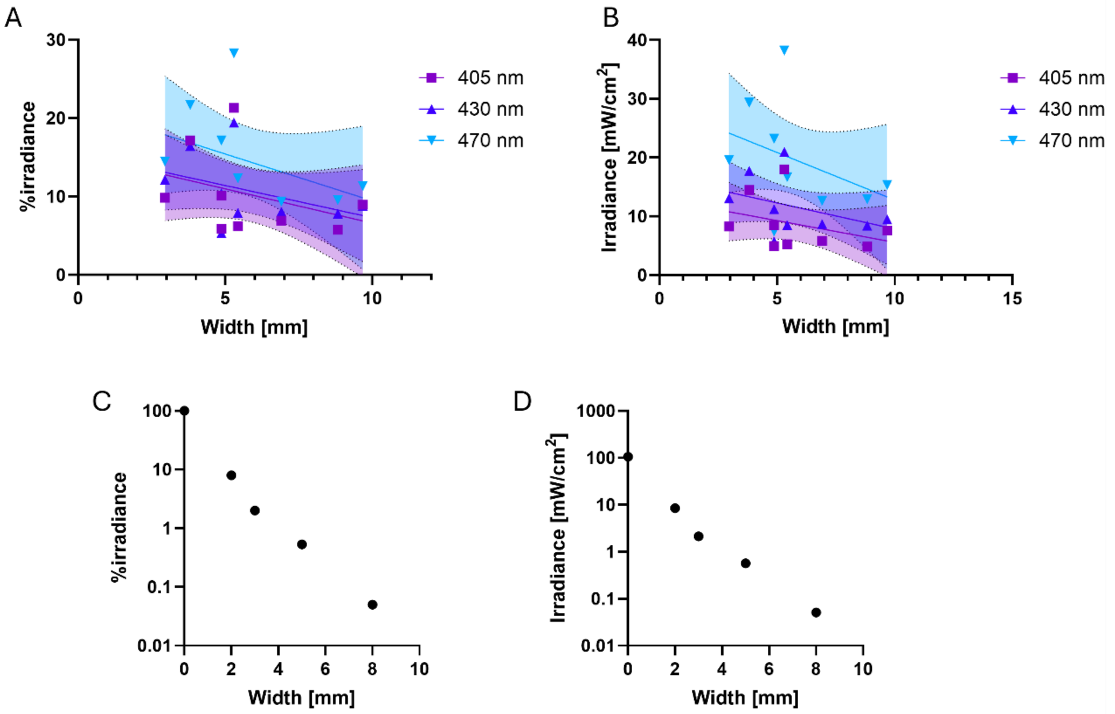

**Figure S15**. Using the setup shown in **Figure S14**, (A,B) irradiance measured after crossing through tumours of different sizes at the indicated wavelengths, and (C,D) irradiance measured after crossing through chicken breast slices of different widths at 470 nm. The %irradiance reflects the fraction of light that remained after passing through the sample, normalised to each LED's unobstructed output.

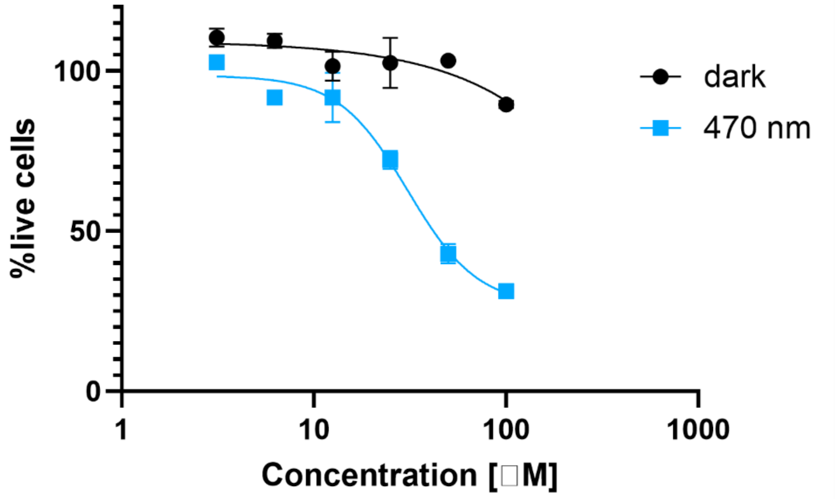

**Figure S16**. Dose-response curves on the viability of MDA-MB-231 cells (MTS assay) of **1C** under dark vs illumination (420 nm, 8 mW/cm$^2$, 40 s) conditions for 72 h, giving an $IC_{50}$ (light) = 30.8 μM.

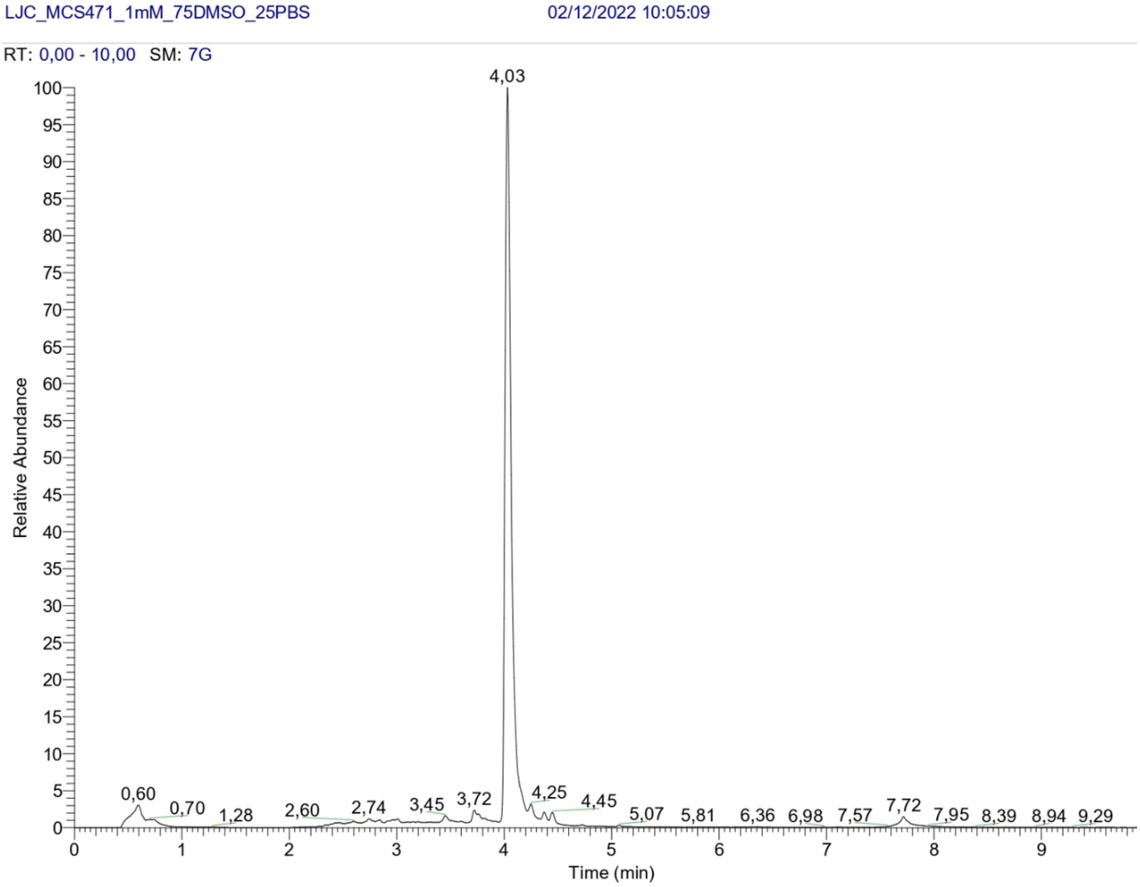

**Figure S17**. HPLC traces of **1H**.

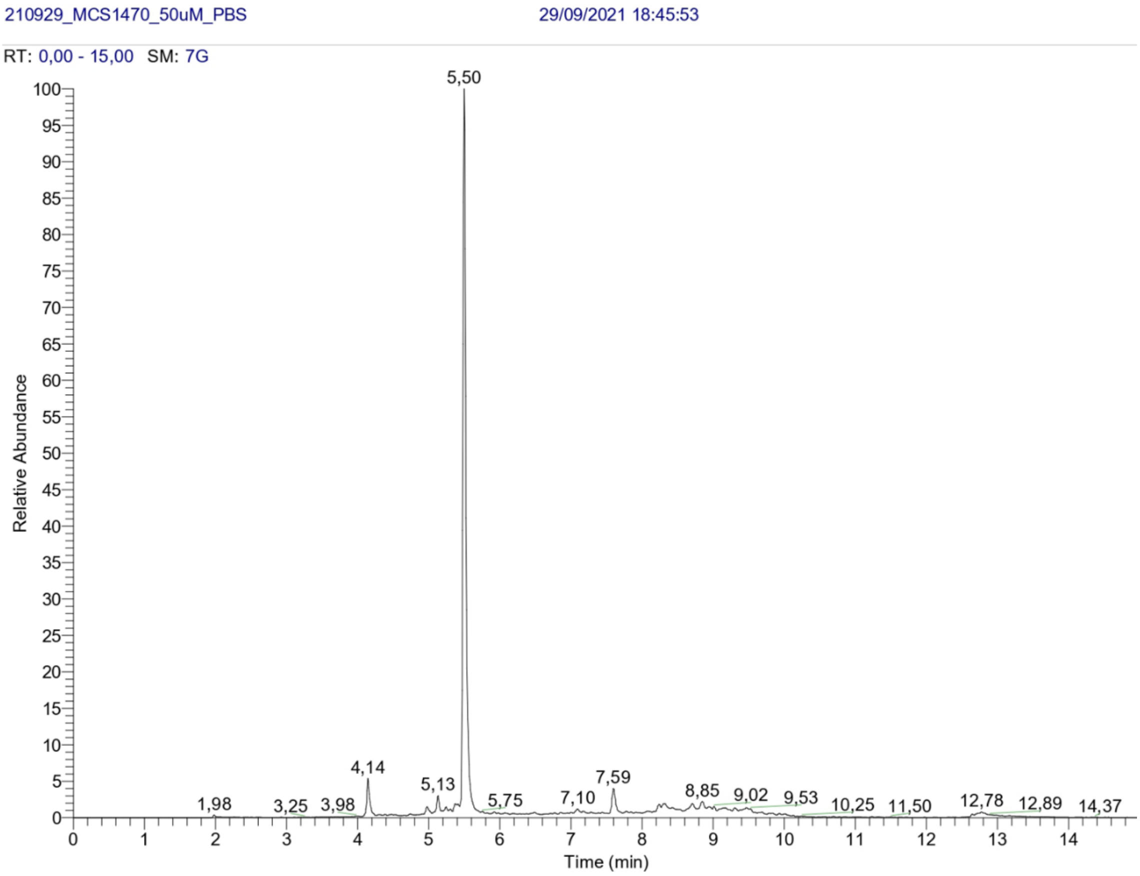

**Figure S18**. HPLC traces of **2C**.

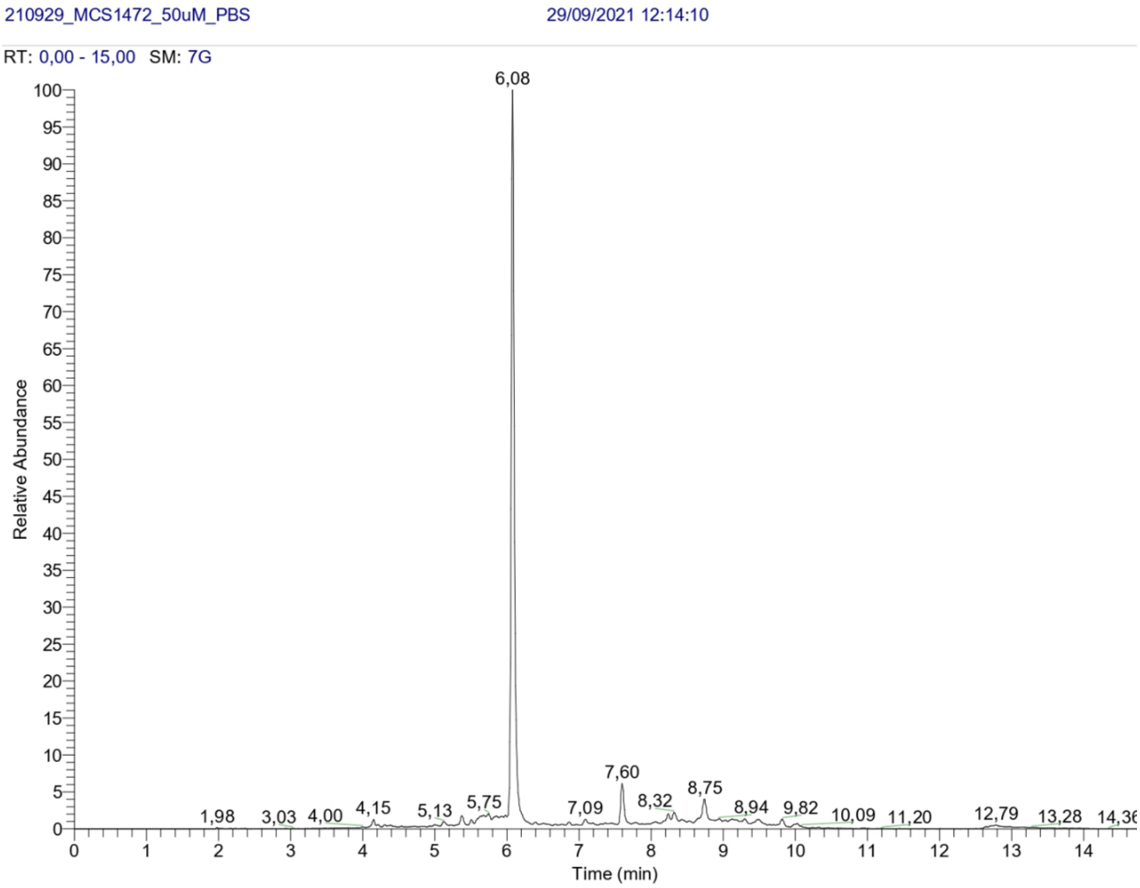

**Figure S19**. HPLC traces of **3C**.

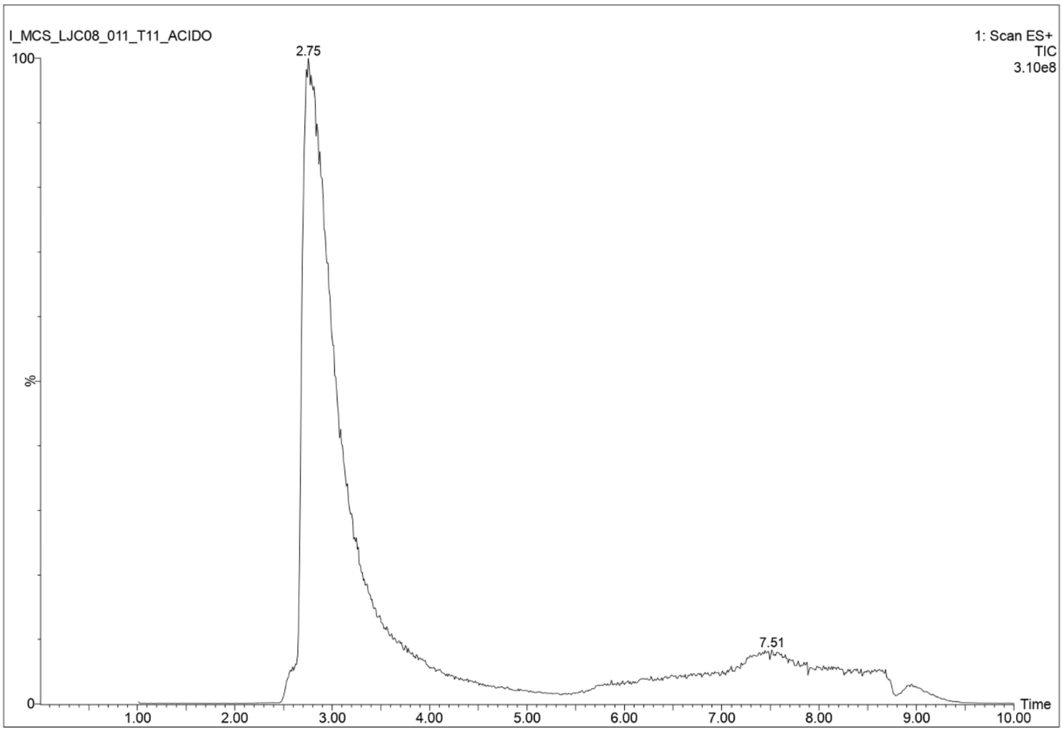

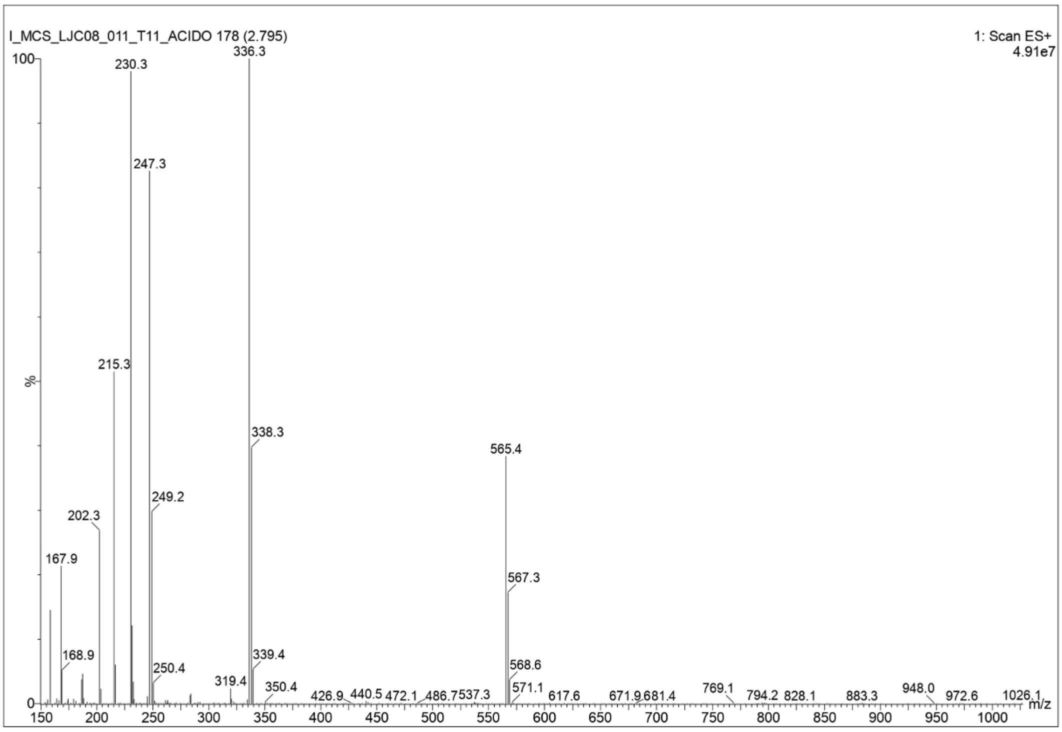

**Figure S20**. HPLC traces of **1H**.

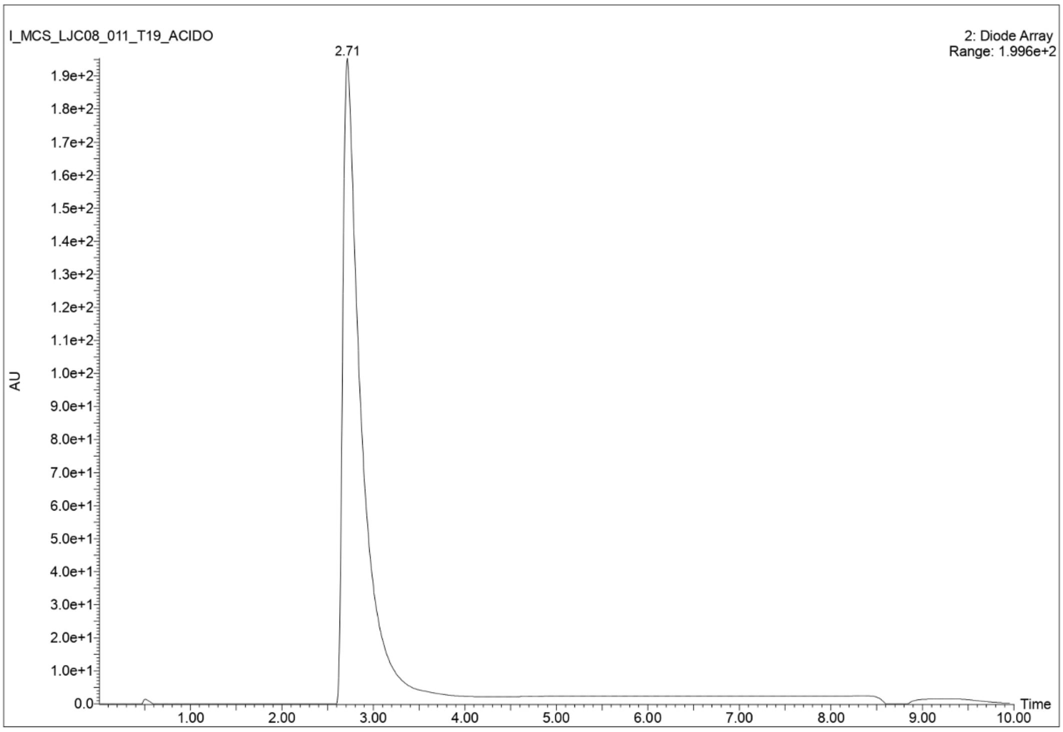

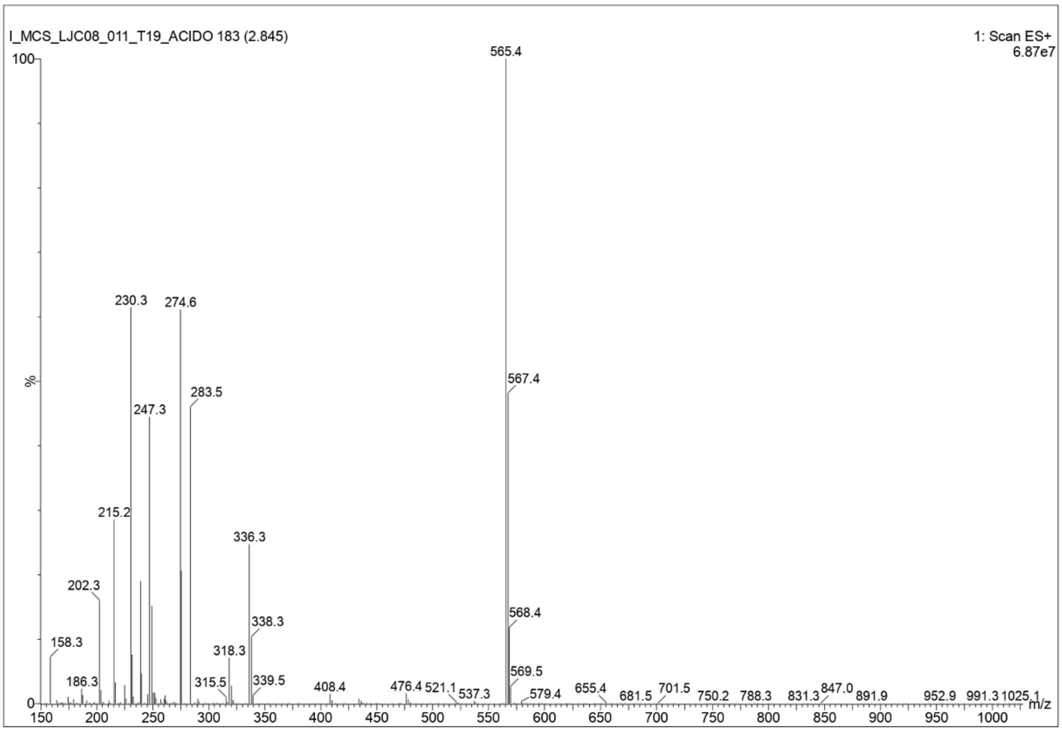

**Figure S21**. HPLC traces of **2H**.

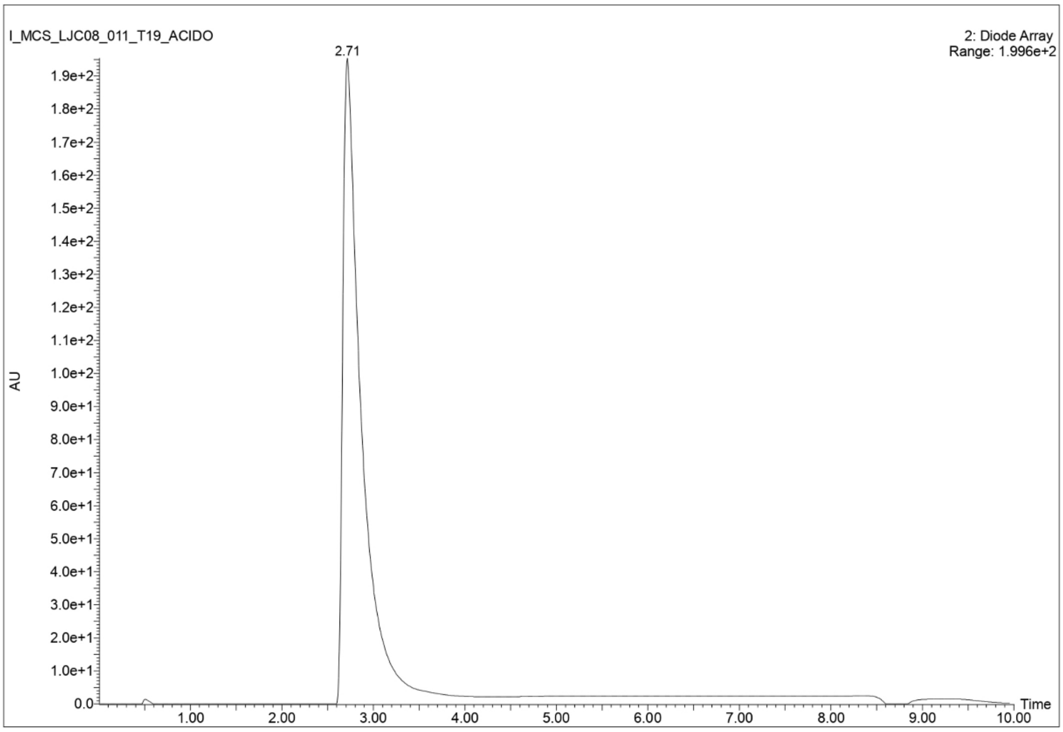

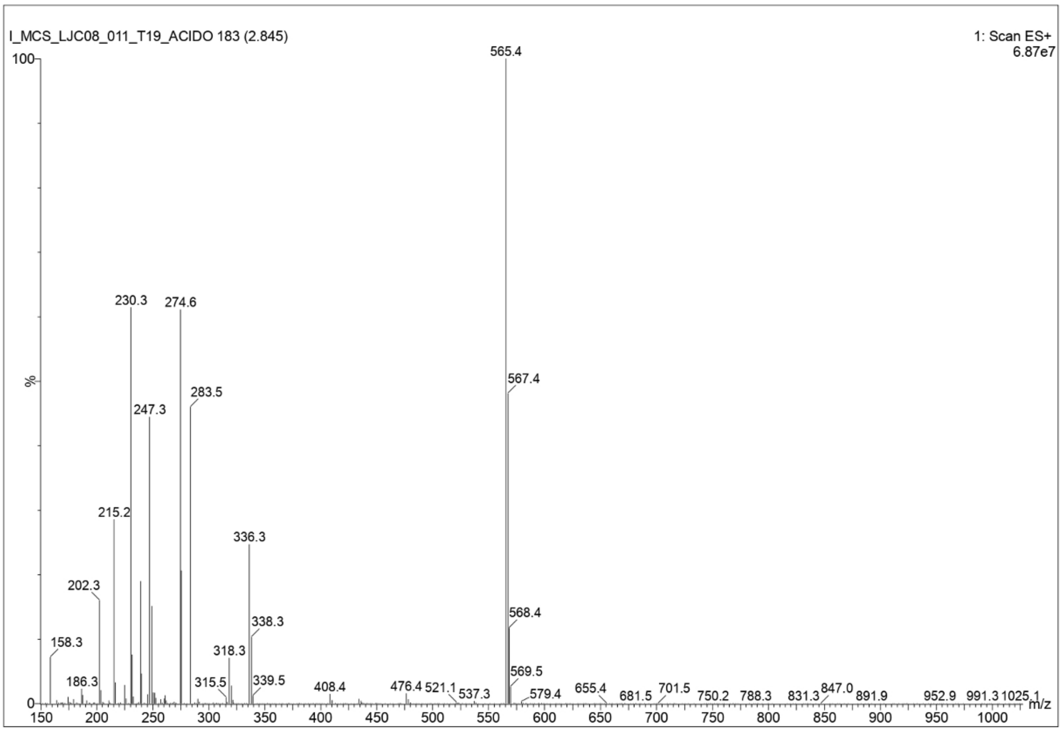

**Figure S22**. HPLC traces of **3H**.

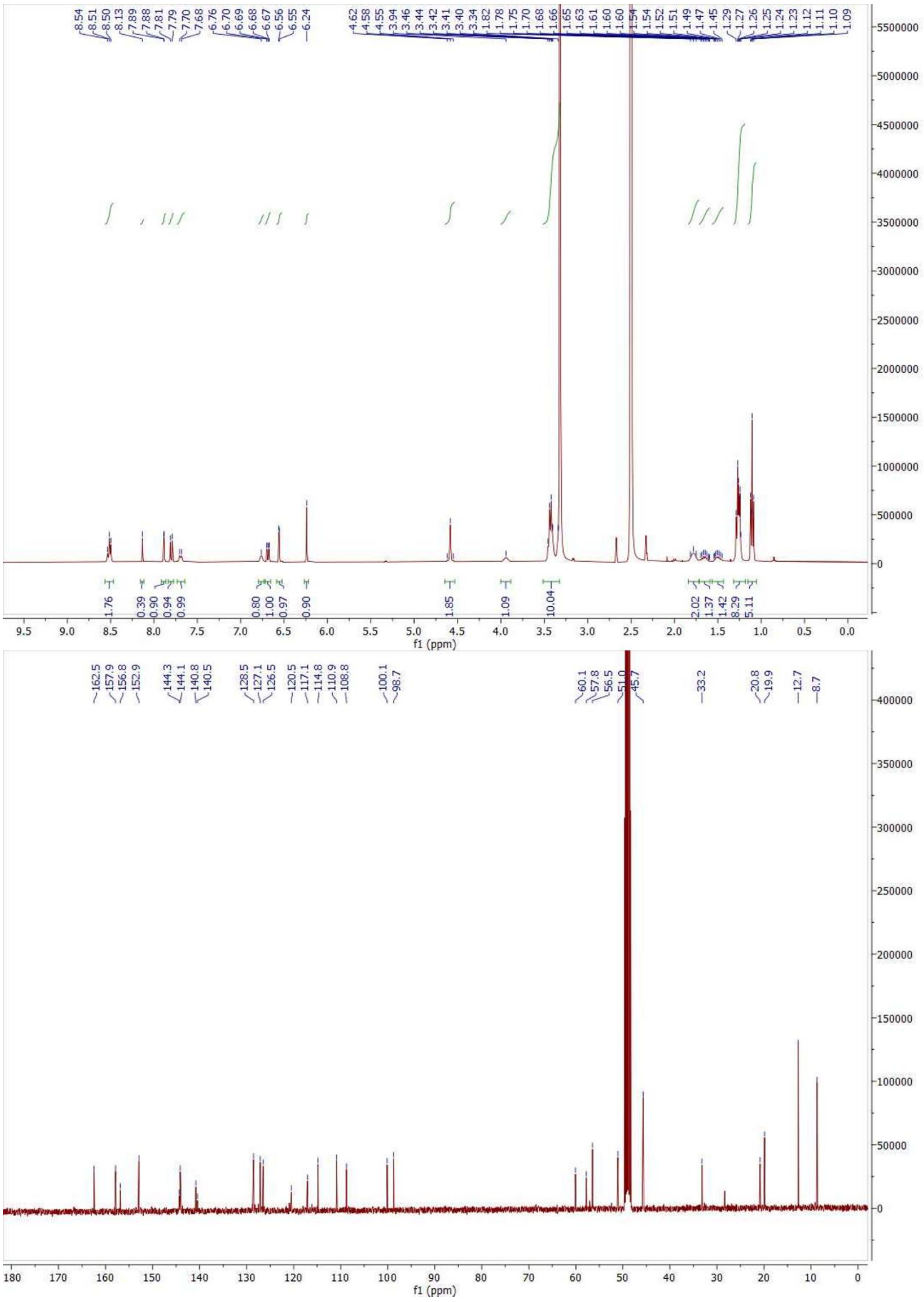

**Figure S23**. $^{1}H$ and $^{13}C$ NMR spectra of **1C**.

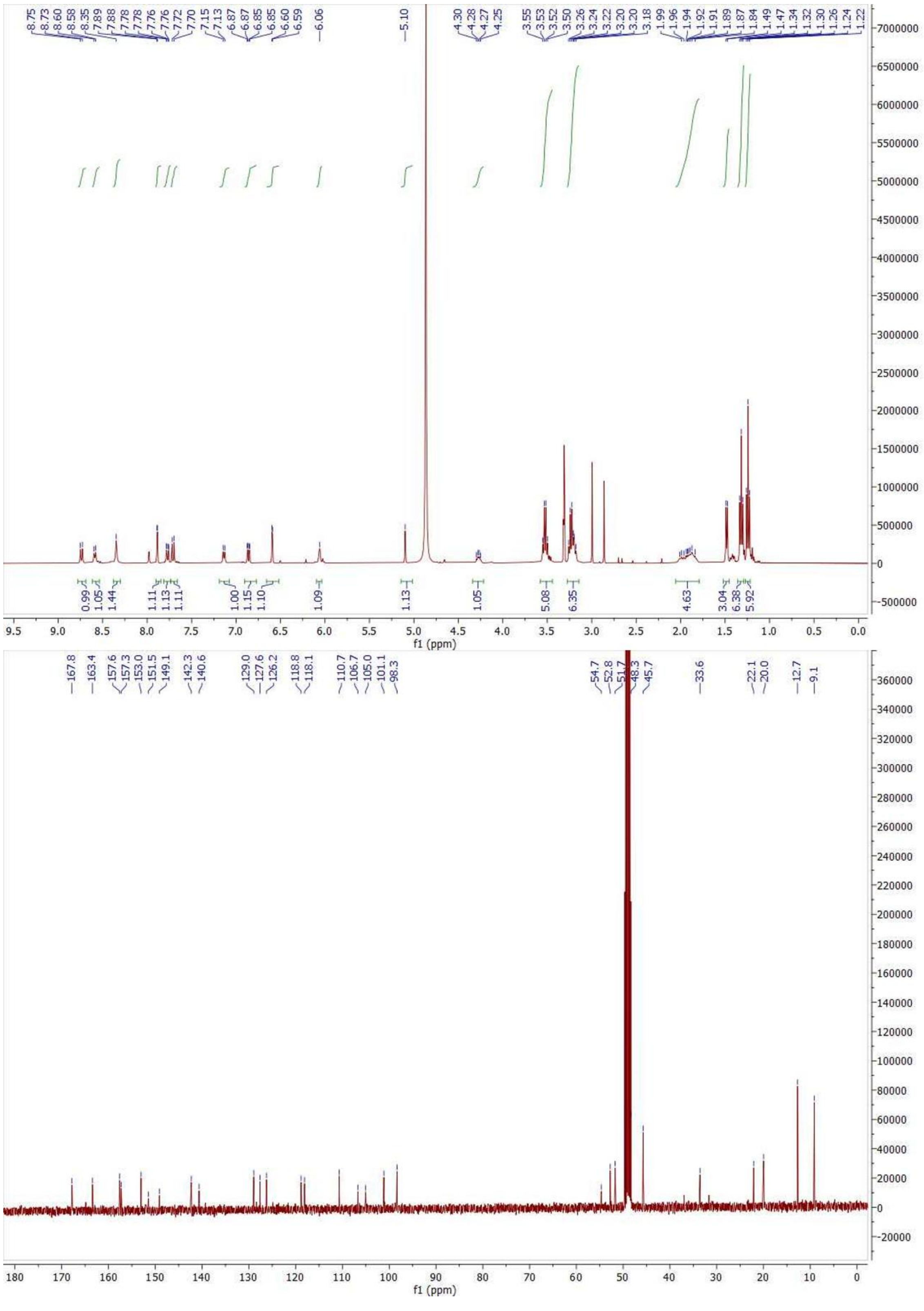

**Figure S24**. $^{1}H$ and $^{13}C$ NMR spectra of **2C**.

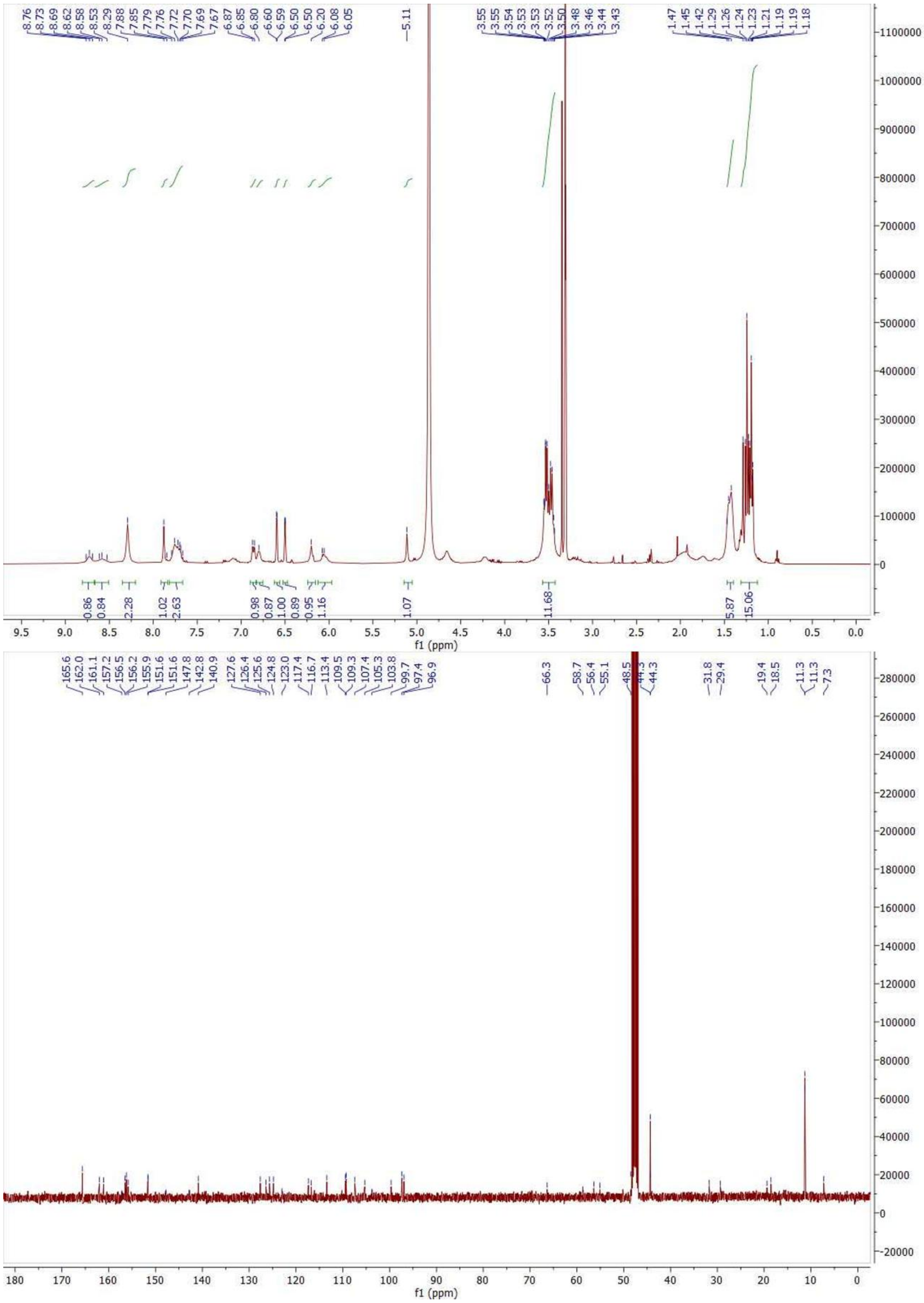

**Figure S25**. $^{1}H$ and $^{13}C$ NMR spectra of **3C**.

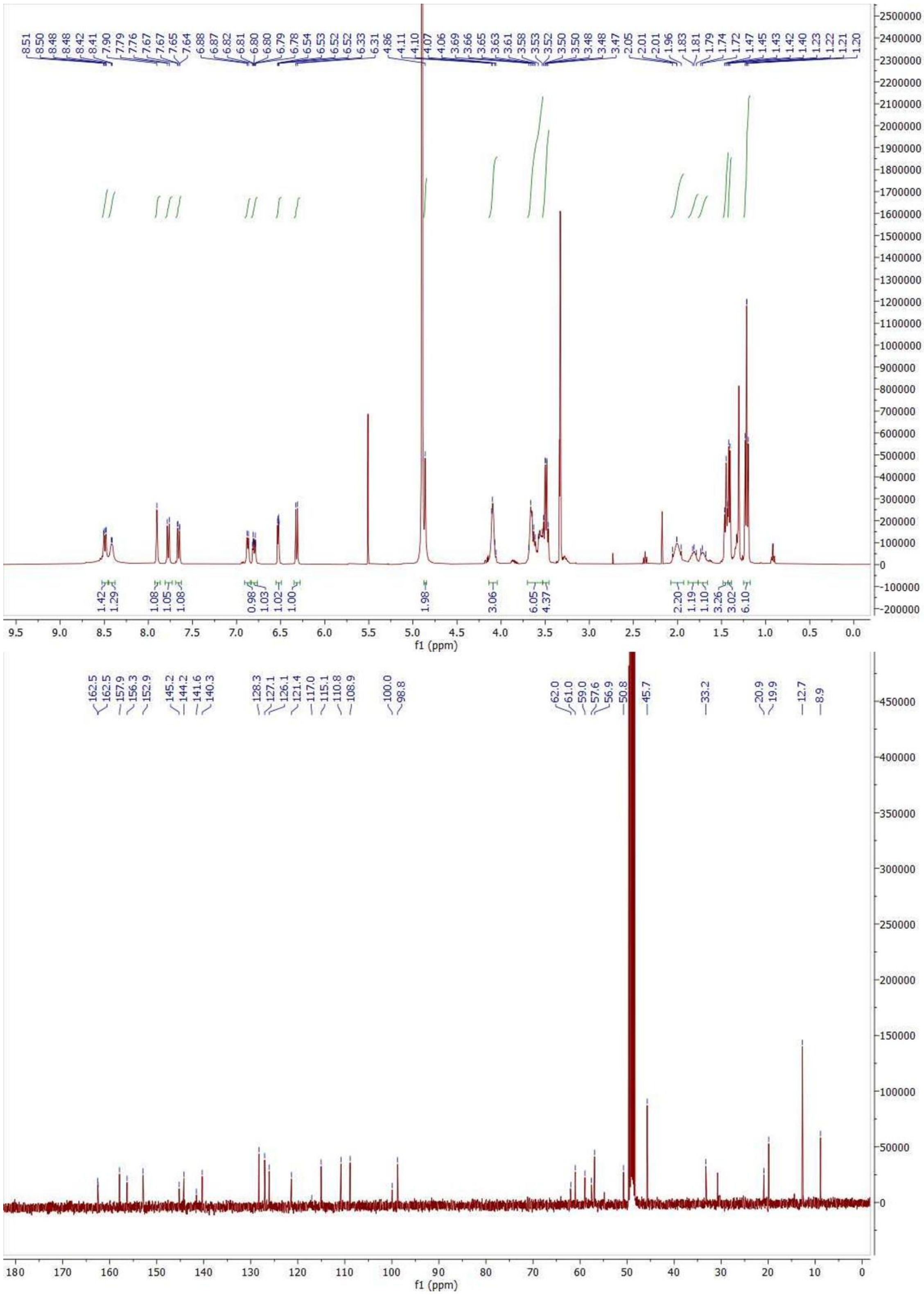

**Figure S26.** $^{1}H$ and $^{13}C$ NMR spectra of **1H**.

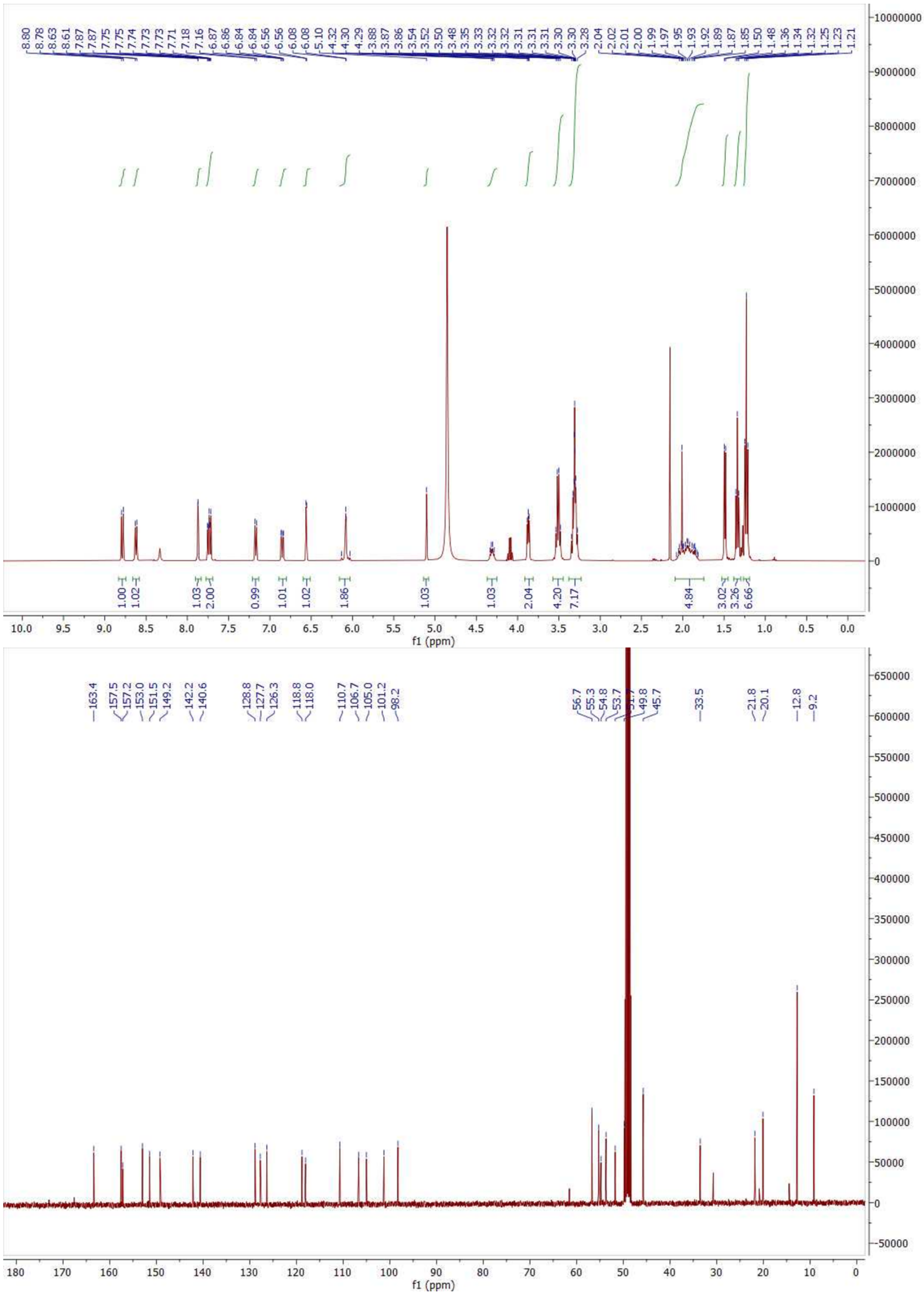

**Figure S27**. $^{1}H$ and $^{13}C$ NMR spectra of **2H**.

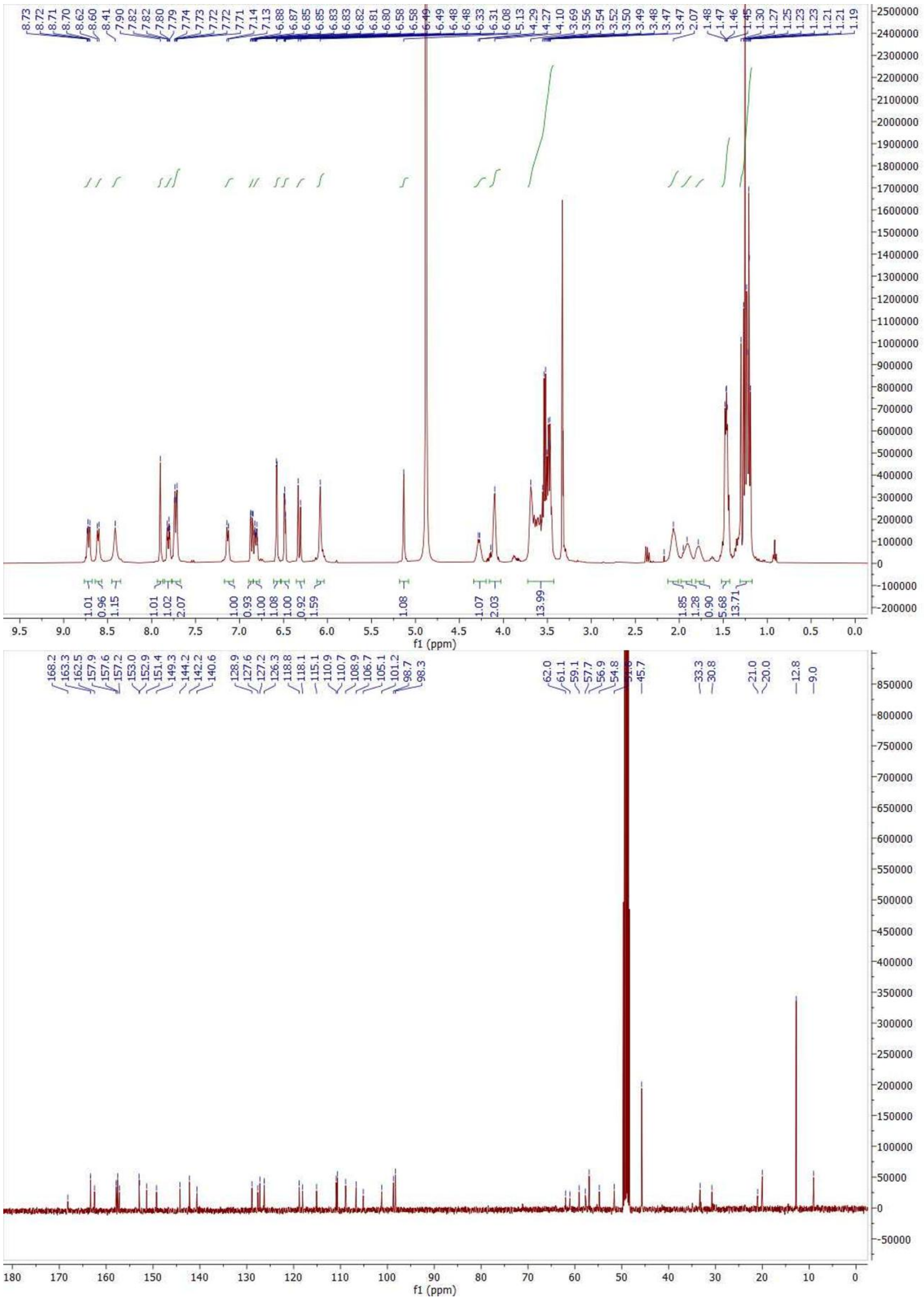

**Figure S28**. $^{1}H$ and $^{13}C$ NMR spectra of **3H**.
